# Epigenomic tomography for probing spatially-defined molecular state in the brain

**DOI:** 10.1101/2022.11.24.517865

**Authors:** Zhengzhi Liu, Chengyu Deng, Zirui Zhou, Ya Xiao, Shan Jiang, Bohan Zhu, Lynette B. Naler, Xiaoting Jia, Danfeng (Daphne) Yao, Chang Lu

**Author notes:** These authors contributed equally (Zhengzhi Liu, Chengyu Deng). Corresponding author (C.L.).

## Abstract

Spatially-resolved genomic profiling is critical for understanding biology in a spatially ordered organ such as mammalian brain. Single-cell spatial genomic assays were developed recently for this purpose but they remain costly and labor-intensive for examining brain tissues across substantial dimensions and surveying a collection of brain samples. Here we demonstrate a new approach, brain epigenomic tomography, that maps spatial epigenomic variations at the scale of centimeters. We profiled neuronal and glial fractions of mouse neocortex slices with 0.5 mm thickness using a low-input technology. H3K27me3 or H3K27ac features across these slices were grouped into clusters based on their spatial variation patterns. Our approach reveals striking dynamics in frontal and caudal cortex due to kainic acid-induced seizure, linked with transmembrane ion transporter, exocytosis of synaptic vesicles, and secretion of neurotransmitter. Epigenomic tomography provides a powerful and cost-effective tool for profiling and discerning brain samples based on their spatial epigenomes.

## Introduction

Profiling genome-wide molecular biology yields rich information on processes critically involved in development and diseases. Genomic, epigenomic and transcriptomic profiles on tissue samples reveal how gene transcription and expression are regulated by different layers of the molecular machinery. During such profiling, the spatial context is often the key information for understanding the tissue-level biology ranging from microenvironment effects to cytoarchitectural functions and organization. Various spatially sensitive approaches have been developed over recent years to probe transcriptomics(Chen et al., 2015; Eng et al., 2019; Lee et al., 2014; Liu et al., 2020; Lubeck et al., 2014; Rodriques et al., 2019; Wang et al., 2018), epigenomics(Deng et al., 2022a; Deng et al., 2022b), genomics(Zhao et al., 2022), and multi-omics(Liu *et al*., 2020; Zhao *et al*., 2022) with resolution up to single cells. These single-cell spatial genomic techniques provide useful insights into the myriad cell types involved and their spatial organization in the tissue.

Among all potential profiling efforts on various tissue types, spatially-resolved profiling of brain epigenomics takes on special significance. Distinct cytoarchitectonic areas in the brain are associated with different functions (Rakic et al., 2009). In addition, epigenomic landscape remains largely plastic during the development and lifespan of a brain cell(Lister et al., 2013) and provides dynamic mediation of both genetic and environmental factors. Spatial brain epigenomics likely yields rich information about normal brain development and neurodevelopmental disorders.

Although sampling a subset of single cells extracted from various brain regions is feasible(Liu et al., 2021), epigenomic profiling of a large area or volume of brain tissue with single-cell resolution remains challenging given the complexity associated with stitching up a huge number of single-cell data sets with spatial identifications, both experimentally and computationally. Single-cell approach may be feasible for creating a reference brain atlas. However, screening and comparing a number of brain samples using this approach will be prohibitive in the foreseeable future due to cost and labor.

Here we demonstrate mapping of epigenomic tomography for characterization of the spatial epigenomic state in the brain. In our approach, mouse neocortex is dissected in a mouse brain slicer to create 0.5 mm-thick neocortex slices that each generates sufficient material for epigenomic profiling of neurons and glia separately, using a low-input technology MOWChIP-seq (Cao et al., 2015; Liu et al., 2022; Zhu et al., 2019). We then create epigenomic tomography separately on neuronal (NeuN+) and glial (NeuN-) fractions by recording the variation in the histone modification signal across the neocortex in the coronal direction. We examine tri-methylation of histone 3 at lysine 27 (H3K27me3) and acetylation of histone 3 at lysine 27 (H3K27ac) which are known repressive and enhancer marks respectively(Zhao and Garcia, 2015) and shown to be highly differentiating of brain regions(Ma et al., 2018). We further apply k-means clustering to group the epigenomic features at specific loci into clusters based on their patterns of variation across the neocortex. Our approach also allows us to compare two brain samples by constructing a double epigenomic tomography. We applied this approach to study the spatially dynamic changes in the mouse neocortex due to induced seizure.

Epigenomic tomography provides a practical and general approach for surveying a large brain area to characterize spatially dynamic processes. This cost-effective approach will potentially allow examination of a sizable collection of brain samples in different conditions to gain fundamental understanding of critical developmental and aberrant processes.

## Results

### Construction of an epigenomic tomography by low-input profiling of brain slices

We dissected the brain from ten-week-old male mice, sliced the neocortex section (roughly 1 cm long) into 19-21 coronal sections with 0.5 mm thickness, and isolated left and right neocortex from each section to produce 38-42 brain slice samples from each neocortex (Fig. 1A). We then homogenized each slice sample and sorted nuclei into neuronal (NeuN+) and glial (NeuN-) fractions after NeuN labeling and fluorescence-activated nuclei sorting (FANS). Neuronal and glial fractions of each slice sample were profiled separately for histone modification using a low-input technology MOWChIP-seq(Cao *et al*., 2015; Zhu *et al*., 2019), generating two replicates using 4,000 nuclei per library on each fraction. We chose to examine H3K27me3 and H3K27ac for their high specificity to brain regions (Ma *et al*., 2018). ChIP-seq data were of high quality (Supplementary Table S1). For example, M4 (Mouse No. 4) and M5 neuronal datasets averagely had 11,069 peaks, 18.4% in fraction of reads in peaks (FRiP), and 0.97 in Pearson’s correlation coefficient between replicates for H3K27me3; 43,951 peaks, 14.4% in FRiP, and 0.90 in correlation between replicates for H3K27ac.

**Figure 1.**
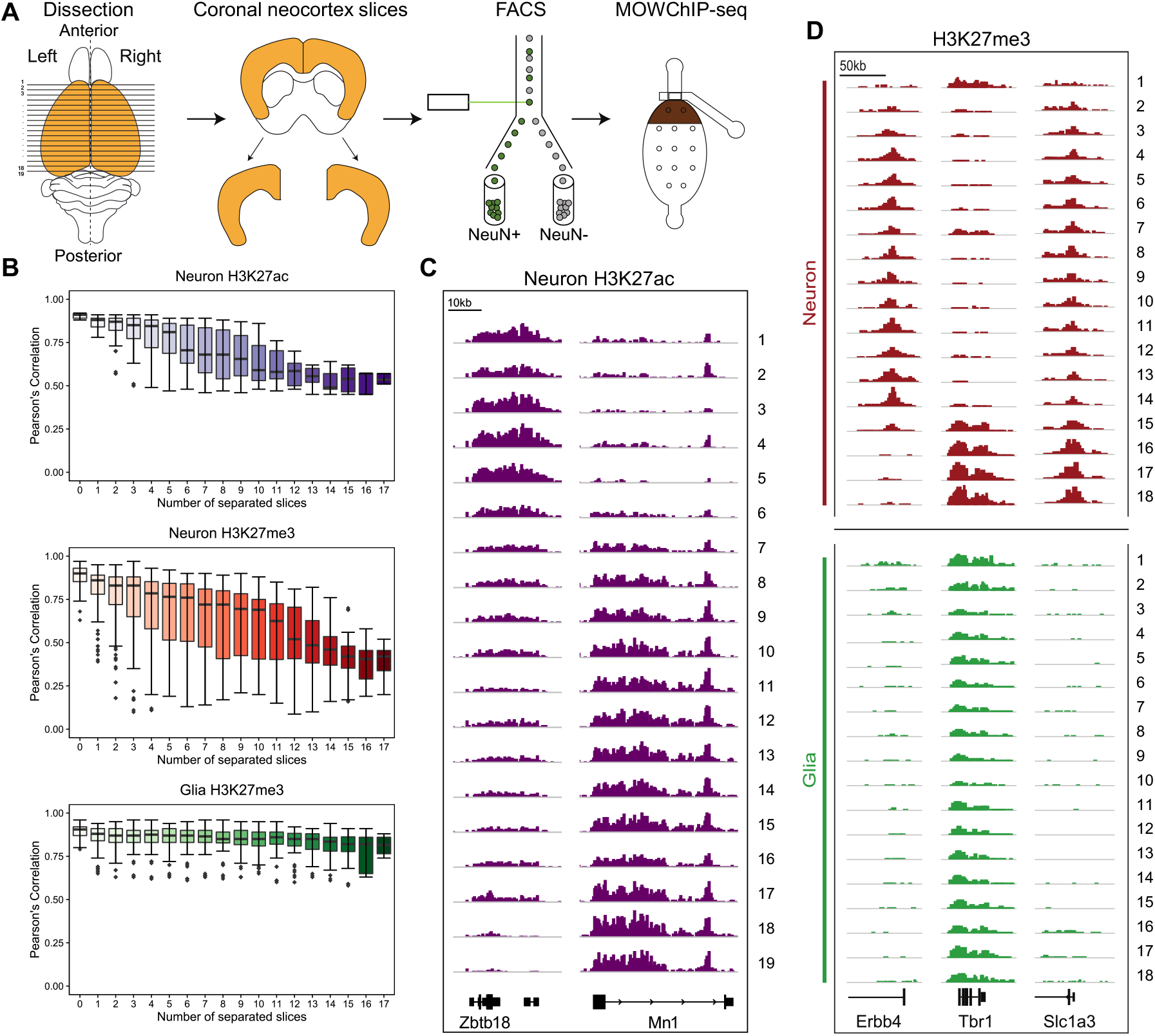
Profiling spatial variations in H3K27me3 and H3K27ac across adult mouse neocortex. (A) A schematic of workflow to create epigenomic tomography. The process includes dissection of mouse neocortex, production of coronal slices of left and right neocortices, nuclei sorting to generate neuronal and glial fractions, and low-input MOWChIP-seq to produce genome-wide histone modification profiles on each slice. (B) Pearson’s correlation between slices that were separated by various number of slices. Horizontal line shows median; box shows interquartile range (IQR); whiskers show ±1.5 × IQR. (C) Normalized H3K27ac signal in mouse neurons (M5). (D) Normalized H3K27me3 signals in neurons and glia from M2.

We calculated Pearson’s correlation coefficient between slices separated by various numbers of slices in their genome-wide histone modification profiles (Fig. 1B). Closely adjacent slices were more strongly correlated than slices that were spaced further apart in the neuronal profiles of both H3K27me3 (average r varying from 0.88 between two adjacent slices to 0.39 with 17 slice separation) and H3K27ac (average r varying from 0.90 between two adjacent slices to 0.54 with 17 slice separation). In contrast, the glial fraction of the slices showed very small variation in the H3K27me3 landscape (with an average r of 0.81 when the slices were separated by 17 slices). Given the high data quality, the various patterns of variation in ChIP-seq signal across the neocortex slices were readily observed at important genes associated with brain functions and activities (Fig. 1C and 1D). *Zbtb18* is involved in brain development and its mutation causes intellectual disability and epilepsy in human(Heng et al., 2022). *Tbr1* regulates neuron identity and *Tbr1* null mice showed severe defects in frontal cortex(Bedogni et al., 2010).

With each slice from the neocortex profiled using our low-input epigenomic technology, we aim to create an epigenomic tomography that captures the spatial variation in the chromatin state and reflects major brain state dynamics. In the case of H3K27me3, we chose high-confidence peaks having overlap with promoter regions for the construction of the tomography. One cell type (i.e. neuron or glia) from each slice has two ChIP-seq datasets/replicates associated with H3K27me3. Top 30% of all peaks by their confidence score in each ChIP-seq dataset were first selected. We then generated a common consensus peak set for all slices by combining the peaks that were present in at least 2 datasets. The consensus peak set was further filtered by removing peaks that did not overlap with promoter regions. Finally, we applied k-means clustering to group consensus peaks with similar pattern of variation across the brain region to generate the epigenomic tomography visualized in the form of heatmap. We produced 4 epigenomic tomographies using data on left and right neocortices of M1 and M2 (Fig. 2A-D). Although the clusters in each tomography varied in the number of consensus peaks included, they showed highly similar patterns of spatial variation across slices. Furthermore, clusters with similar patterns show highly overlapped gene ontology (GO) terms despite that the 4 tomographies were constructed using different data. Specifically, clusters I and IV are consistently linked with gene expression regulation, biosynthetic processes and development in all samples. Cluster II is associated with development, morphogenesis, and proliferation. Cluster V is linked with chemical stimulus and sensory perception, with the only exception in M2 left. In spite of the similarity in their spatial pattern and associated GO terms, the corresponding clusters in various neocortex samples contained different numbers of peaks. However, these peaks had a substantial level of overlap (Fig. 2E). For example, cluster I contained 555, 419, 540, 594 peaks in M1 left, M1 right, M2 left and M2 right neocortex samples, respectively, with 156 common peaks presented in the cluster I of all these samples.

**Figure 2.**
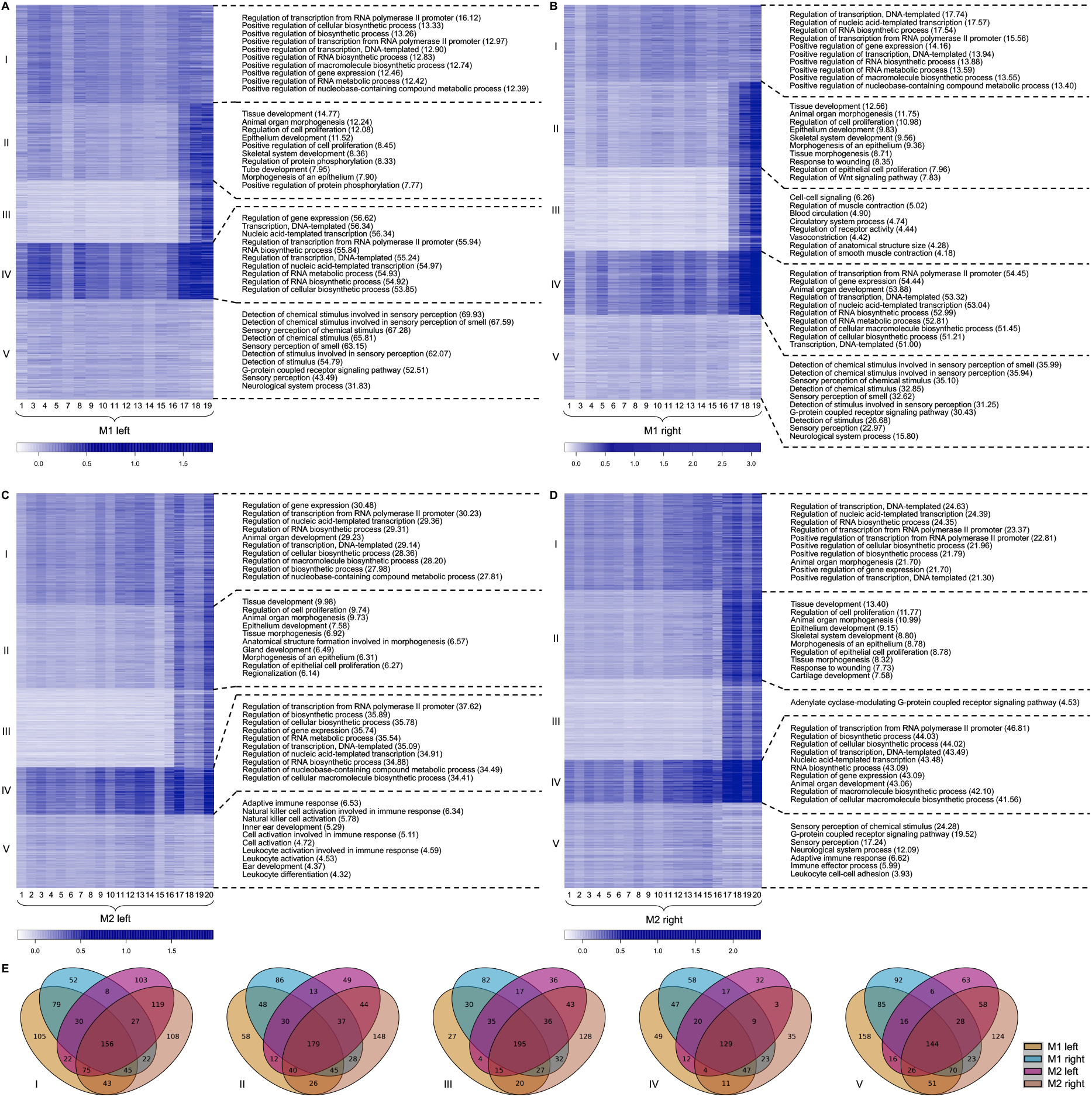
Epigenomic tomography generated by k-means clustering of promoter H3K27me3 peaks using neuronal data on neocortex halves and top GO biological process terms associated with each cluster. (A) Neuronal fraction of M1 left neocortex; (B) Neuronal fraction of M1 right neocortex; (C) Neuronal fraction of M2 left neocortex; and (D) Neuronal fraction of M2 right neocortex; (E) Overlap of peaks among clusters of the same spatial pattern from all four epigenomic tomographies. The heatmaps (A-D) show the intensity of consensus promoter H3K27me3 peaks across slices 1-19/20 grouped by K-means clustering. Top 10 GO biological process terms were shown if the total number of GO terms is greater than 10. The full list of GO terms is presented in Supplementary Dataset S1.

We decided that the epigenomic tomography approach was the most useful in the comparison of two or more brain samples. We combined two sets of H3K27me3 data (M1 left and M1 right, a “double tomography”) and conducted k-means clustering to compare the two halves of the neocortex from the same mouse (Fig. 3A). We show that the left and right neocortices present high level of similarity in their pattern of variation across slices (Fig. 3A). The consensus peaks were grouped into 5 clusters with different patterns of spatial variation. Most clusters present distinct and very related GO terms. For example, Cluster I is associated with development; Cluster II is closely related to development and morphogenesis; Cluster IV is related to transcription, RNA biosynthetic process and development; Cluster V is linked to chemical stimulus and sensory processes. Furthermore, the processing of spatial epigenomic data on a different mouse (M2 left and M2 right) yielded very similar clusters in terms of variation patterns. The enriched GO terms are also highly similar, especially for Clusters I, II, IV and V (Fig. 3B). The grouped peaks in the same spatial cluster had substantial overlap between the two epigenomic tomographies constructed using data from two mice (Fig. 3C). Finally, the neocortices on the same side of M1 and M2 brains also showed high similarity in their spatial clustering (Supplementary Fig. S1). The epigenomic tomographies yielded by k-means clustering of M1 left-M2 left and M1 right-M2 right data (Supplementary Fig. S1) were also highly consistent with the ones produced by different combination of the data sets (Fig. 3) in terms of their spatial pattern and the enriched GO terms. Interestingly, 3 out of 4 “double tomographies” (Fig. 3B and Supplementary Fig. S1 A-B) showed enriched term “G-protein coupled receptor signaling pathway” for Cluster III. Taken together, our results suggest that the epigenomic tomography approach creates fairly reproducible profiles when neocortices from different sides or different control mice were examined.

**Figure 3.**
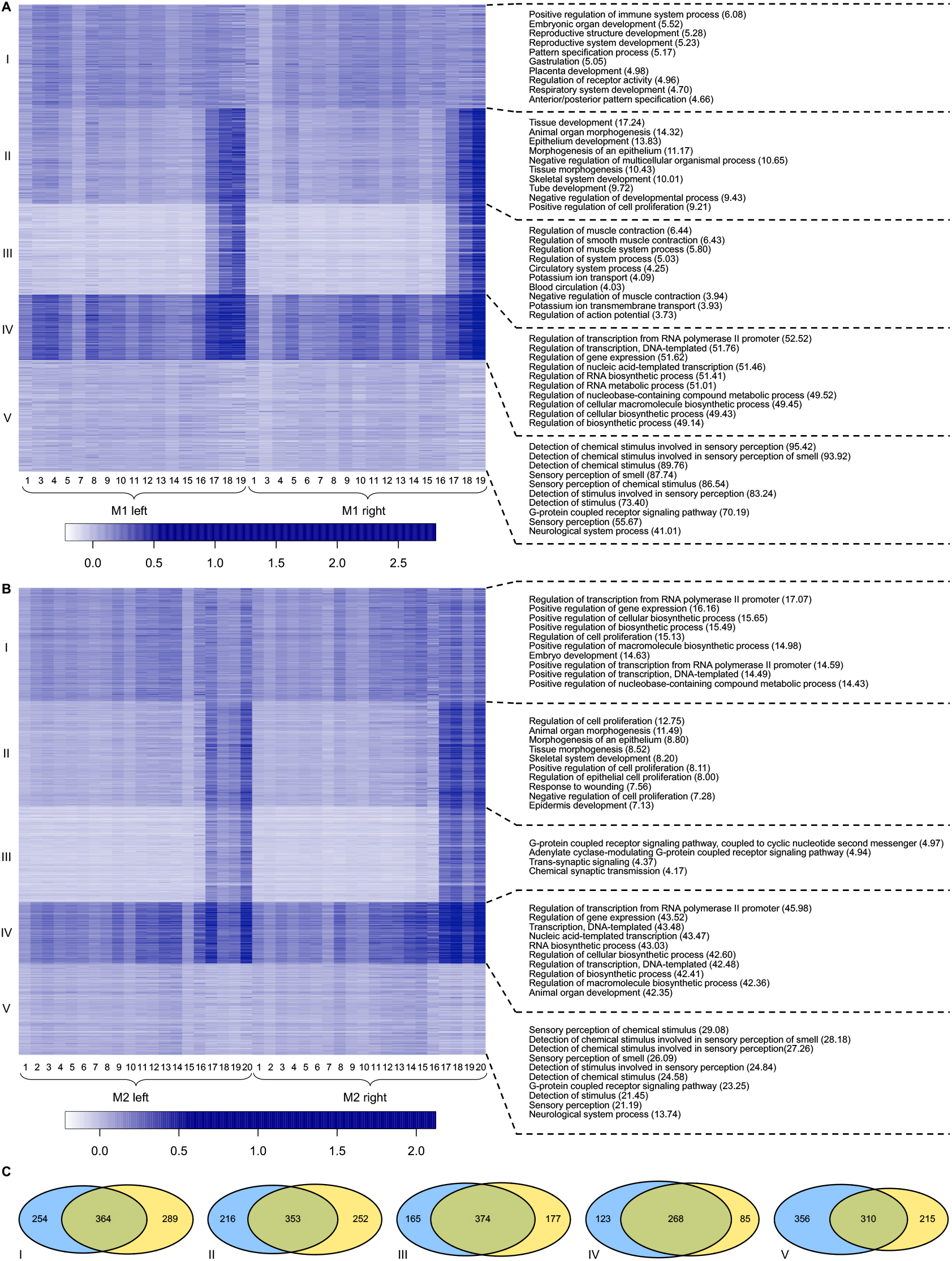
Double epigenomic tomography generated by k-means clustering of promoter H3K27me3 peaks using neuronal data on both left and right neocortices and top GO biological process terms associated with each cluster. (A) Neuronal fraction of both left and right neocortices of M1. (B) Neuronal fraction of both left and right neocortices of M2. (C) Overlap of peaks between clusters of the same spatial pattern from the two epigenomic tomographies. The heatmaps (A-B) show the intensity of consensus promoter H3K27me3 peaks across slices 1-19/20 of two brain samples grouped by K-means clustering. Top 10 GO biological process terms were shown if the total number of GO terms is greater than 10. The full list of GO terms is presented in Supplementary Dataset S2.

In contrast, epigenomic tomographies on glial data show highly similar pattern of spatial variation across the slices among various clusters (Fig. 4 A-B). Thus the peaks were mostly grouped by k-means clustering based on their similarity in the peak intensity. Nevertheless, there remains substantial overlap among peaks of the corresponding clusters from different tomographies (Fig. 4C). Using data on both sides of M1 and M2 neocortices, different combination of the glial data sets yielded similar glial epigenomic tomographies (Fig. 4 and Supplementary Fig. S2).

**Figure 4.**
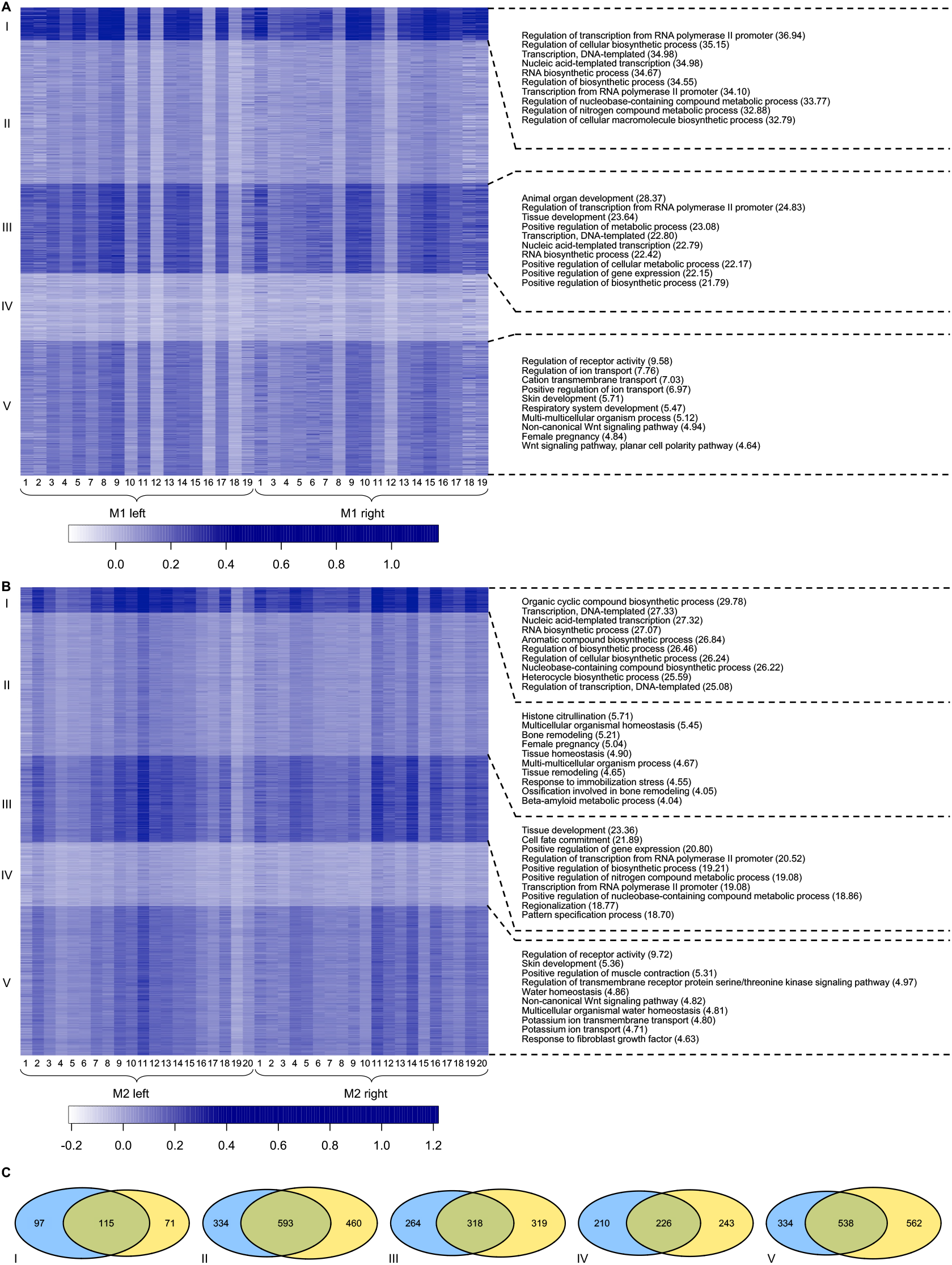
Double epigenomic tomography generated by k-means clustering of promoter H3K27me3 peaks using glial data on both left and right neocortices and top GO biological process terms associated with each cluster. (A) Glial fraction of both left and right neocortices of M1. (B) Glial fraction of both left and right neocortices of M2. (C) Overlap of peaks among clusters of the same spatial pattern from the two epigenomic tomographies. Top 10 GO biological process terms were shown if the total number of GO terms is greater than 10. The full list of GO terms is presented in Supplementary Dataset S3.

We explored the link between our epigenomic tomography with mRNA tomography. We applied SMART-seq2 to profile neuronal fraction of the right neocortex slices. We computed the RNA signal over the genebody of all RefSeq mouse genes and the H3K27me3 signal over the promoter regions of the same genes. Pearson’s correlation between RNA signal and promoter H3K27me3 signal across all slices was then calculated to evaluate correlation between the two marks for a particular gene. Among the 4,443 genes that passed the quality filter, the majority (2,839, or 64%) had negative correlation between RNA and H3K27me3 while 495 genes showed high negative correlation (r < −0.5) (Supplementary Fig. S3A). For example, *Tanc2* and *Zic1* are associated with signaling during brain development (Guo et al., 2019; Inoue et al., 2007; Kim et al., 2021) (Supplementary Fig. S3B).

### Epigenomic tomography reveals spatially-specific changes in the neocortex due to seizure

We examined whether the epigenomic tomography reflects molecular dynamics in the epigenomic landscape of the brain due to a major pathological change. We induced seizures in mice via kainic acid injection(Levesque and Avoli, 2013; Sharma et al., 2018). Electroencephalographic (EEG) measurement showed that, after a systematic injection of kainic acid into rats, epileptiform activity first starts in the hippocampus and then propagates to the amygdala, the thalamus, and the prefrontal cortex (Ben-Ari et al., 1981; Levesque et al., 2009; Medvedev et al., 2000). Kainate-induced seizure may also cause dendritic injury in the mouse neocortex (Zeng et al., 2007). However, spatial epigenomic changes within the neocortex have not been investigated previously. In our experiment, we examined a mouse injected with kainic acid that exhibited induced seizure (M4) and a control mouse injected with saline (M5). Both mice were euthanized and dissected after 3 d from the time of the injection. We focused on the neuronal fraction from slices in the right neocortex and profiled both H3K27me3 and H3K27ac, with the latter being a mark for active enhancer.

We conducted k-means clustering of H3K27ac peaks that did not overlap with promoters (enhancer H3K27ac peaks) and promoter H3K27me3 peaks, using both seizure and control data in each case. We clustered the enhancer peaks into 6 clusters of distinct patterns and identified 4 clusters that differed substantially between the control and seizure mice (Fig. 5). We roughly divided the mouse neocortex into three parts: frontal cortex (CxF), medial cortex (CxM), and caudal cortex (CxCA). We discovered that the majority of the enhancer differences between seizure and control brains occurred in the CxF (I, III, IV and V) and CxCA (III, V) parts. The GO terms associated with these spatially differential clusters (I, III, IV, and V) were highly concentrated on exocytosis of synaptic vesicles, secretion and transport of neurotransmitter, and regulation of transmembrane ion transporter activity. Ion transportation across membrane and exocytosis form the basis of electrical and chemical signaling in nerve cells (The principles of nerve cell communication, 1997). This agrees with the fact that seizure is associated with severely perturbed neural activity characterized by significant changes to extracellular and intracellular ion concentrations and increased neuronal firing (Gonzalez et al., 2019; Raimondo et al., 2015). Interestingly, Ras protein signal transduction was involved in one of the spatially differential clusters (V). Ras protein is also known as Ras GTPase, which belongs to the group of small GTPases(Lu et al., 2016). One of the negative regulators of Ras protein, *SYNGAP1* (Keller et al., 2017), causes epilepsy and intellectual disability with its mutation(Berryer et al., 2013). Mutations of *SYNGAP1* were detected in epileptic encephalopathies, a condition where constant epilepsy leads to worsening brain function(Carvill et al., 2013). *Syngap1* KO mice also showed lower threshold to induced seizures(Clement et al., 2012). We examined the H3K27ac landscape around *Syngap1* and observed stronger signal in CxF slices and weaker signal in CxCA slices for the seizure sample than the control (Supplementary Fig. S4). Interestingly, locomotory behavior was also identified as a GO term associated with cluster IV. Seizure impacts locomotory behavior, likely through disrupting suprachiasmatic nucleus activity(Möller et al., 2019; Murphy and Burnham, 2003; Stewart and Leung, 2003). We examined closely the peaks associated with the GO term locomotory behavior and discovered that the top 2 genes with the most associated cluster IV peaks were *Grin2a* and *Gnao1. Grin2a* mutation causes brain development delay, language difficulties and epilepsy in human(Lesca et al., 2013; Strehlow et al., 2019; Venkateswaran et al., 2014). *Gnao1* mutation causes epilepsy, development delay and movement disorder(Feng et al., 2018; Nakamura et al., 2013). The enhancer peaks associated with the genes (identified by examining the correlation between the peak alternation and the gene expression) were substantially altered due to seizure (Supplementary Fig. S5-6), suggesting that epigenomic tomography is effective for picking up important perturbed brain processes. Other genes ranked in the top 20 in terms of their number of associated cluster IV peaks included *Negr1*(Singh et al., 2018), *Apba2*(Lin et al., 2019), *Atp2b2*(Smits et al., 2019), *Lsamp*(Catania et al., 2008), *Mapt*(Shaw-Smith et al., 2006; Strang et al., 2019), *Cacna1c*(Nyegaard et al., 2010), *Dab1*(Feng and Cooper, 2009), *Fgf14*(Hoxha et al., 2019), *Cacnb4*(Tadmouri et al., 2012), *Cntn2*(Stogmann et al., 2013), and *Lrrtm1*(Francks et al., 2007), which all play significant roles in brain development and functions.

**Figure 5.**
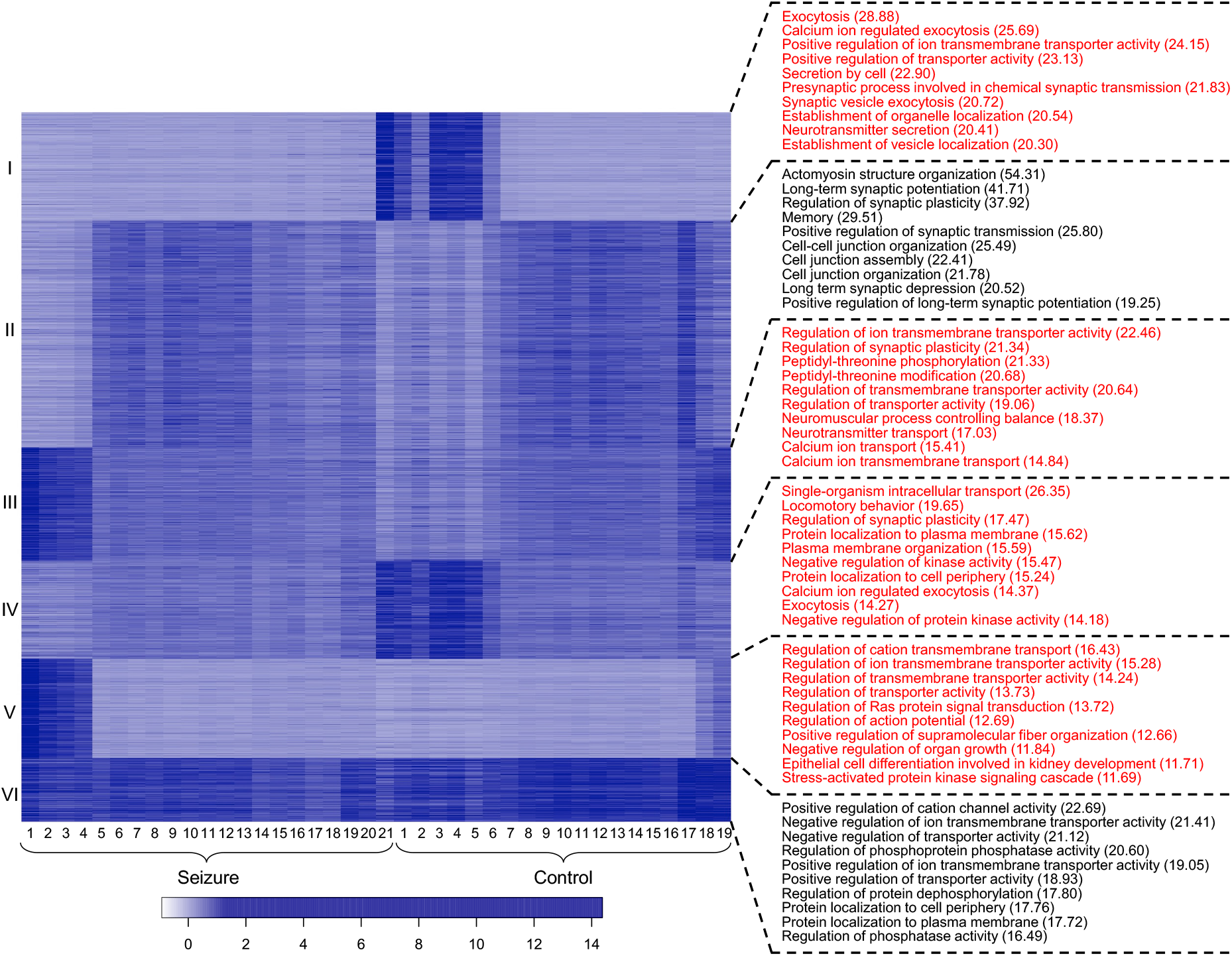
Double epigenomic tomography generated by k-means clustering of enhancer peaks using neuronal data on the right neocortex of both seizure and control mice and top GO biological process terms associated with each cluster. k=6. Top 10 GO biological process terms were shown if the total number of GO terms is greater than 10. GO terms associated with differential clusters are in red. The full list of GO terms is presented in Supplementary Dataset S4.

We examined the effect of k value used in k-means clustering on detection of spatially differential peaks between seizure and control brains. The elbow method suggested a range for the optimal k value (Supplementary Fig. S7). We showed that k-means clustering with k values in the optimal range (4-7) generally yielded similar results in terms of the spatially differential peaks between seizure and control and the GO terms associated with the spatially differential clusters (Fig. 6). With increased k value (i.e. the number of clusters), the total number of spatially differential peaks increased from 14,778 to 20,068 and 21,860 when k increased from 4 to 5 and 6, respectively. Further increase to k of 7 slightly reduced the total number of spatially differential peaks (from 21,860 to 20,506 when k increased from 6 to 7). There was a high degree of overlap among the spatially differential peaks discovered using various k values, with 14,634 peaks identified by every k-means clustering analysis with k = 4-7.

**Figure 6.**
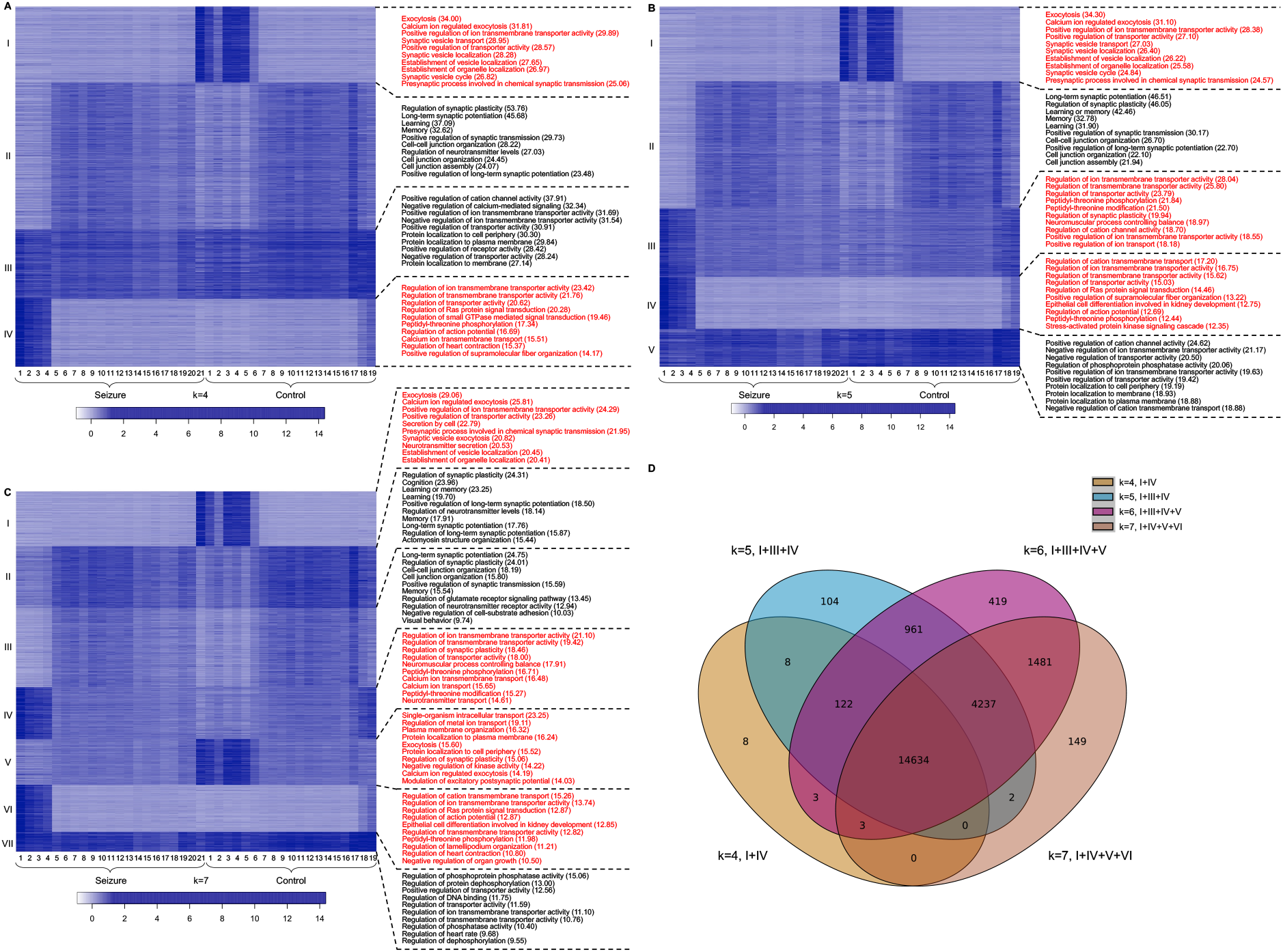
The effect of k value on the double epigenomic tomography of seizure/control. k=4 (A); 5 (B); 7 (C). (D) Overlap of the spatially differential enhancer peaks generated at various k values. Top 10 GO biological process terms were shown if the total number of GO terms is greater than 10. GO terms associated with differential clusters are in red. The full list of GO terms is presented in Supplementary Dataset S5.

Promoter H3K27me3 tomography also revealed difference between control and seizure neocortices (Supplementary Fig. S8). Clusters V and VI showed significant alternation in the spatial variation pattern between the two cases. The changes were also mostly located in the CxF part of the neocortex, in agreement with what we observed in the enhancer (H3K27ac) tomography. However, each of the two clusters did not yield significant GO terms. When the two clusters were combined, “GTPase mediated signal transduction” and “Ras protein signal transduction” terms were found to be enriched, in agreement with the GO terms identified from the enhancer tomography. We did not discover GO terms related to transmembrane ion transport as with the enhancer tomography, suggesting that each histone mark is linked with different molecular processes.

## Discussion

We generated epigenomic tomography by profiling genome-wide histone modifications in neuronal or glial cells from about 20 slices of adult mouse neocortex using low-input ChIP-seq. We observed that the mouse neocortex could be divided into three distinct blocks (frontal, medial, and caudal) based on the spatial variation in H3K27me3 or H3K27ac of neurons. The histone modification signal tends to be fairly uniform within these blocks but abrupt change in the signal intensity may occur at their interfaces. Such division was not observed in glial profiles. We constructed the epigenomic tomography using k-means clustering to group peaks with similar spatial variation pattern into the same clusters. Interestingly, some of these clusters showed strong enrichment in similar GO terms, suggesting that epigenomic features associated with highly related molecular processes and functions tend to exhibit similar spatial variation pattern.

Clustering of epigenomic tomographic data from two samples (i.e. double tomography) allows comparison of spatial variation pattern for the same peaks across brain samples. We examined H3K27me3 and H3K27ac tomographies in the neuronal fraction of mouse neocortices that underwent injection of kainic acid or saline to investigate the effect of induced seizure to epigenomic tomography. We identified significant differences between the two conditions, primarily in the frontal and caudal sections of the neocortex. The spatially differential peaks between control and seizure revealed a number of molecular processes that were closely involved in seizure. Both enhancer H3K27ac tomography and promoter H3K27me3 tomography suggested the importance of GTPase-based signal transduction in seizure induction. In addition, the enhancer tomography also revealed the involvement of transmembrane ion transporter and exocytosis in seizure, which is in agreement with existing knowledge(Gonzalez *et al*., 2019; Raimondo *et al*., 2015).

In our analysis of epigenomic tomography, GO analysis was used to discover the biological processes, cellular components, and molecular functions implicated by the clusters of spatially differential peaks. We envision that this can be a general approach for extracting biological insights from epigenomic tomography comparison. Although GO analysis is susceptible to misuse and misinterpretation(Yon Rhee et al., 2008), such analysis of k-means clustered spatially differential peaks may generate quick references to altered brain state and varied molecular processes.

There is space for further improvement of epigenomic tomography in terms of its spatial resolution and cell-type specificity. Because our MOWChIP-seq technology allows profiling using as few as 100 cells (compared to 4,000 nuclei per assay used in this study), use of thinner slices and separation into more fractions should be feasible in principle. Slices as thin as 10-20 μm can be produced using a cryostat and the nuclei can be sorted into 4 or more cell types based on molecular markers(Hauberg et al., 2020).

Our epigenomic tomography is conceptually similar to “RNA tomography” performed using low-input RNA-seq(Junker et al., 2014). Compared to the focus on individual genes or loci in RNA tomography, our use of K-means clustering provides a way to visualize the spatial dynamics and group similarly trending peaks to yield direct insights into relevant biological processes.

Compared to the state-of-the-art single-cell spatial omic technologies, epigenomic tomography reveals changes across a much larger area at a much lower cost, albeit with a coarser spatial resolution. Epigenomic tomography does not reveal information on microenvironment, cell-cell communication or cell type composition at the level of current single-cell spatial technologies. Nevertheless, epigenomic tomography presents genomewide dynamics across different regions of the brain and potentially at the organ level. Furthermore, although it is potentially feasible to create a reference brain atlas with singlecell spatial resolution, such approach is currently too expensive and labor-intensive for study and comparison of a sizable collection of brain samples. Our low-cost tomography approach can fill the gap if the changes or dynamics occur across a substantial physical dimension. We envision that our epigenomic tomography approach will facilitate discovery of new epigenomic mechanisms underlying developmental or disease conditions in brain that are spatially sensitive.

## Methods

### Mouse brain tissues

All procedures were conducted in accordance with NIH guidelines and were approved by the Institutional Animal Care and Use Committee (IACUC) at Virginia Tech. Adult (P56) C57BL/6J male mice were purchased from the Jackson Laboratory and housed on a 12 h light/dark cycle at 23 °C with food and water ad libitum for 2 weeks before experiments.

M1-3 were euthanized using compressed CO_2_ followed by cervical dislocation. Mouse brain was then isolated and cut into coronal slices with a thickness of 0.5 mm using an adult mouse brain slicer (BSMAS005-1, Zivic Instruments) on ice. Left and right neocortex sections were dissected from each slice and immediately frozen in dry ice. The dissected neocortex slice samples (a total of 18-21 slices for either left or right neocortex) were stored at −80 °C until use.

M4 and M5 were treated for the study of kainic acid-induced seizure. 2 mg/ml kainic acid (K0133, LKT Laboratories) in sterile saline was prepared freshly. M4 was injected intraperitoneally with repeated 5 mg/kg doses of kainic acid at 30 min intervals until behavioral seizures were observed (after 2 doses)(Puttachary et al., 2015) (Supplementary Video S1). M5 (control) was injected with the same amount of sterile saline at 30 min intervals for 2 doses. M4 and M5 were euthanized by cervical dislocation 3 d after the injections and death was confirmed by bilateral thoracotomy. Mouse brain was processed to extract 0.5 mm-thick left and right coronal neocortex slices and the slice samples were stored at −80 °C until use.

### Nuclei extraction and sorting

Nuclei extraction from dissected brain tissue was performed using previously described protocol(de la Fuente Revenga et al., 2021; Jiang et al., 2008) with minor modifications. All steps were conducted on ice and centrifugation was conducted at 4 °C. A neocortex slice was thawed on ice and transferred to a glass tissue homogenizer (Sigma, d9036-1SET). 3 ml of ice-cold nuclei extraction buffer was added [0.32 M sucrose, 5mM CaCl_2_, 2 mM Mg(Ac)_2_, 0.1 mM EDTA, 10 mM tris-HCl, 0.1% (v/v) Triton X-100, freshly added 1% protease inhibitor cocktail (PIC, Sigma-Aldrich), 0.1 mM PMSF, 1 mM DTT, and 0.06 U/μl ribonuclease (RNase) inhibitor (2313A, Takara Bio, needed when mRNA extraction was performed)]. The tissue was homogenized with 10 times of slow douncing using a loose pestle and 15 times of douncing using a tight pestle (D9063, Sigma-Aldrich). The homogenate was filtered through a 40-μm cell strainer (22363547, Fisher Scientific) into a 15-ml centrifuge tube and centrifuged at 1000 rcf for 10 min at 4 °C. Supernatant was discarded and the pellet was resuspended in 0.5 ml of ice-cold nuclei extraction buffer. 0.75 ml of 50% (w/v) iodixanol (made by mixing 4 ml of OptiPrepTM gradient (Sigma-Aldrich) and 0.8 ml of diluent [150 mM KCl, 30 mM MgCl2, and 120 mM tris-HCl]) was added. The suspension was gently mixed by inverting the tube up and down 5 times and then centrifuged at 10,000 rcf for 20 min. Supernatant was carefully removed and the pellet incubated with 100 μl of 2% (w/v) normal goat serum (50062Z, Life Technologies) in Dulbecco’s PBS (DPBS, Life technologies) on ice. The pellet was then resuspended and transferred into a new Eppendorf tube with an addition of 200 μl of 2% normal goat serum. To differentiate neurons from glia, 10 μl of 2 ng/μl Anti-NeuN antibody conjugated with Alexa 488 (MAB377X, EMD Millipore) was mixed with the suspension. The mixture was incubated at 4 °C for 1 h on a rotator. The nuclei solution was sorted using FACS (BS FACSAria, BD Bioscience) to separate neuronal (NeuN+) and glial (NeuN-) fractions.

### Chromatin fragmentation

#### M1-2

4,000 nuclei per replicate was used so that both neuronal and glial fractions of every neocortex slice sample yielded enough material for two ChIP-seq replicates. The nuclei suspension (containing 8,000 nuclei) was topped up to 1 ml using Dulbecco’s PBS (DPBS; 14190144, Life Technologies). 200 μl of 1.8M sucrose, 10 μl of 1M CaCl_2_, and 3 μl of 1M Mg(Ac)_2_ were then added sequentially. The sample was mixed gently by inverting the tube up and down and then incubated for 15 min on ice. The suspension was centrifuged at 3,000 rpm for 15 min at 4 T. Supernatant was carefully removed, and the pellet was resuspended in 10 μl DPBS. 0.1 μl of PIC, 0.1 μl of 100 mM PMSF and 10 μl of lysis buffer [4% Triton X-100, 100 mM tris (pH 7.5), 100 mM NaCl, and 30 mM MgCl_2_] were added, mixed by vortexing and then incubated at room temperature for 10 min. 1 μl of 0.1 M CaCl_2_ and 0.4 μl of 100 U/μl MNase (88216, Thermo Fisher Scientific) were then added, mixed by vortexing and incubated at room temperature for 10 min. 2.22 μl of 0.5 M EDTA (pH 8) was added to quench the rection. The sample was vortexed and incubated on ice for 10 min before it was centrifuged at 16,100 g for 5min. The supernatant (~20 μl, containing chromatin) was transferred to a new tube and 60 μl of IP buffer [20 mM Tris-HCl, 140 mM NaCl, 1 mM EDTA, 0.5 mM EGTA, 0.1% (w/v) sodium deoxycholate, 0.1% SDS, 1% (v/v) Triton X-100] was added to obtain a total volume of ~80 μl. The solution was finally split for two ChIP-seq replicates. The chromatin was stored on ice before loaded into the MOWChIP device.

#### M4-5

200 μl of sorted neuronal nuclei suspension from one neocortex slice sample was used to prepare chromatin (containing ~30,000 nuclei). 2 μl of PIC, 2 μl of 100 mM PMSF and 200 μl of lysis buffer were added, mixed by vortexing and incubated at room temperature for 10 min. 4 μl of 1 M CaCl_2_ and 5 μl of 100 U MNase (88216, Thermo Fisher Scientific) were added, mixed by vortexing and incubated at room temperature for 10 min. 44 μl of 0.5 M EDTA was then added, mixed by vortexing and incubated on ice for 10 min. The mixture was finally centrifuged at 16,100g for 5 min at 4 °C. Supernatant containing fragmented chromatin (~ 440 μl) was collected into a new 1.5 ml tube and placed on ice before use. 70 μl of chromatin (containing chromatin from ~4,000 nuclei) was used for each MOWChIP-seq assay for consistency with data on M1-2.

### Antibody coating on IP beads

5 μl of Dynabeads Protein A (10002D, Thermo Fisher Scientific) were used in each MOWChIP assay. Beads were washed twice with 150 μl of IP buffer [20 mM Tris-HCl, pH 8.0, 140 mM NaCl, 1 mM EDTA, 0.5 mM EGTA, 0.1% (w/v) sodium deoxycholate, 0.1% SDS, 1% (v/v) Triton X-100] and then resuspended in 150 μl of IP buffer. An antibody that targets a specific modified histone was added to coat the beads: anti-H3K27ac (ab4729, Abcam) at 0.125 μg/assay or anti-H3K27me3 (39155, Active Motif) at 0.5 μg/assay. The bead/antibody suspension was incubated on a rotator at 4 °C overnight. Beads were then washed with IP buffer twice and resuspended in 5 μl of IP buffer for use in MOWChIP.

### MOWChIP

The MOWChIP process, including device fabrication and operation, was performed following our published protocol.(Zhu *et al*., 2019)

### Construction of ChIP-seq libraries

~4,000 neuronal or glial nuclei were used to produce each ChIP-seq library and two technical replicates were generated on each neocortex slice sample. The input libraries were created using the same starting quantity of nuclei (~ 4,000 per library) with the MOWChIP step skipped. Libraries were constructed using 2S Plus DNA Library kit (10009877, IDT) and 2S CDI Adapter S1 indexing kit (10009895, IDT) following the manufacturer’s instructions. 5% (v/v) EvaGreen (31000, Biotium) was added to the amplification mixture to monitor the progress of library amplification. Amplification was stopped once an increase of 2,000 RFU was reached. Library was eluted to 10 μl of low EDTA TE buffer.

### mRNA-seq

mRNA libraries were prepared following Smart-seq2 protocol(Picelli et al., 2014b) with minor modifications. ~7,500 neuronal nuclei (from M3) were used to generate each RNA-seq library, and each slice sample has two technical replicates. 50 μl of neuronal suspension containing about 7,500 nuclei was used for RNA extraction using RNeasy Mini kit (QIAGEN) and RNase-Free DNase Set (79254, QIAGEN). Extracted RNA was concentrated using ethanol precipitation and resuspended in 5 μl of RNase-free water. 4.6 μl of RNA solution, 2 μl of 10 μM oligo-dT primer and 2 μl of 10 mM dNTP mix were mixed, incubated at 72°C for 3 mins and then stored on ice. 11.4 μl of reverse-transcription mix (made from 1 μl of SuperScript II reverse transcriptase (200 U/μl), 0.5 μL of RNase inhibitor (40 U/μl), 4 μL of Superscript II first-strand buffer, 1 μl of DTT (100mM), 4 μl of 5 M Betaine, 0.12 μl of 1 M MgCl_2_, 0.2 μl of TSO (10 μM), 0.58 μl of nuclease-free water) was added to the RNA solution and incubated at 42°C for 90 min, followed by 10 cycles of (50°C for 2 min, 42°C for 2 min), and at 70°C for 15 min. 25 μl KAPA HiFi HotStart ReadyMix, 0.5 μl of 10 μM IS PCR primers, 2.5 μl of 20x Evagreen, and 4 μl of RNase-free water were then added. cDNA was amplified by incubation at 98°C for 1 min, followed by 9-11 cycles (to achieve an increase of 50-100 RFU) of (98°C for 15 s, 67°C for 30 s, 72°C for 6 min). After amplification, cDNA was purified using 0.8× SPRIselect beads. Tn5 transposase (14-17 mg/ml) was produced following a published protocol(Picelli et al., 2014a). Tn5 transposase and T5/T7 transposons were assembled to form Tn5 transposome using a published protocol(Amini et al., 2014). Briefly, 1.5 μl of 100 μM T5 transposon (IDT) was mixed with 1.5 μl of 100 μM pMENTS (a 5’-phosphorylated 19-bp mosaic end complementary oligonucleotide, IDT), while 1.5 μl of 100 μM T7 transposon (IDT) was mixed with 1.5 μl of 100 μM pMENTS. 3 μl of T5-pMENTS solution and 3 μl of T7-pMENTS solution were separately incubated at 95°C for 5 min followed by a slow ramp to 25°C at −0.1°C/sec. 2.5 μl of T5-pMENTS solution was then mixed with 2.5 μl of Tn5 transposase (14-17 mg/ml), while 2.5 μl of T7-pMENTS solution was mixed with 2.5 μl of Tn5 transposase. T5-Tn5 solution and T7-Tn5 solution were separately incubated at 37°C for 1 h to form T5-Tn5 transposome and T7-Tn5 transposome, respectively. 4 μl of T5-Tn5 transposome and 4 μl of T7-Tn5 transposome were mixed to form 8 μl of assembled Tn5 transposome before it was mixed with 1 μl cDNA (containing ~500 pg cDNA) and 1 μl reaction buffer (TNP92110, Lucigen). The reaction mixture was incubated at 37°C for 1 h. 1 μl of 10× stop solution (TNP92110, Lucigen) was added to the reaction mixture to stop tagmentation. Two rounds of 1× SPRIselect beads cleanup were performed to purify tagmented cDNA. The tagmented cDNA was then eluted in 10 μl low EDTA TE buffer. 25 μl KAPA HiFi HotStart ReadyMix was activated at 98 °C for 30 sec and then mixed with 9.5 μl tagmented cDNA and 10 μl Nuclease-free water, followed by incubation at 72 °C for 5 min. 1.5 μl of 25 μM P5 primer, 1.5 μl of 25 μM P7 primer and 2.5 μl 20x Evagreen were then added. The following amplification program was used to achieve a 300-400 RFU increase: 98 °C for 30 s; 11-12 cycles of (98 °C for 10 s, 63 °C for 30 s, 72 °C for 30 s) and 72 °C for 1 min. Amplified library was purified with 0.8× SPRIselect beads and eluted in 8 μl low EDTA TE buffer.

### Library quality control and sequencing

Library size and concentration were measured by Tapestation (Agilent) and KAPA Library Quantification Kit (07960140001, Roche), respectively. Libraries were pooled for sequencing by Illumina HiSeq 4000 with single-end 50 nt reads or NovaSeq 6000 in S4 200 mode with paired-end 100 nt reads. Averagely, 19 million reads were generated for each ChIP-seq library, 12 million reads for each ChIP-seq input library, and 16 million reads for each RNA-seq library.

### ChIP-seq data analysis

Raw FASTQ files were trimmed and QC filtered using Trim Galore! (0.4.1) with default settings. Filtered reads were mapped to reference genome mm10 with bowtie (1.1.2). Uniquely mapped reads were filtered to remove known blacklisted regions using samtools (1.3.1)(Li et al., 2009) and bedtools (2.29.2)(Quinlan and Hall, 2010). H3K27ac peaks were called using MACS2 (2.1.1.20160309) with default settings. H3K27me3 peaks were called using epic2 (0.0.41) with the following settings: -fs 250 --gaps-allowed 4 -fdr 1e-20. Uniquely mapped ChIP and input reads were then extended by 100 bp on both ends. The reference genome was divided into 100-bp bins and signal within each bin was counted for ChIP and input samples. Normalized ChIP-seq signal per bin was calculated as follows:

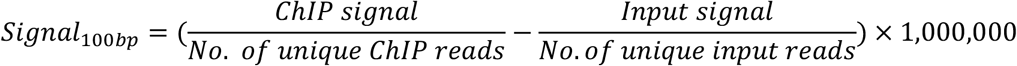

Normalized signal was visualized as genome browser tracks in IGV (2.11.2). The enhancer peak was identified by selecting H3K27ac peak within ± 1M bp from TSS of a gene with the highest correlation with the gene expression (mRNA-seq) data(de la Fuente Revenga *et al*., 2021; Gorkin et al., 2020).

### Construction of epigenomic tomography

Epigenomic tomography was constructed using ChIP-seq datasets from one brain sample or two brain samples (i.e. double epigenomic tomography). Peaks from all datasets of interest were ranked by confidence score (generated by MACS2 or epic2). The top 30% peaks of each dataset were loaded into DiffBind (3.14) and peaks present in at least 2 datasets were selected into a consensus peak set. Consensus peaks were then checked for overlap with promoter regions, defined as ±2 kb from transcription start sites, by bedtools *intersect* function. We selected H3K27me3 peaks that overlap with promoter regions (promoter H3K27me3) and H3K27ac peaks that do not intersect with promoters (enhancers) (Cao *et al*., 2015) when assembling the epigenomic tomography on H3K27me3 and H3K27ac, respectively. We calculated the average normalized read counts in each peak region (i.e. peak intensity) between two replicate ChIP-seq datasets for each slice using deepTools *multiBigwigSummary* (3.5.0). We then created a matrix with the row vectors being consensus peak intensities across all slices under investigation and the column vectors being the intensities across all selected consensus peaks in individual slices. K-means clustering was done with the *kmeans* function of R *stats* package (3.6.2) with the default Hartigan algorithm (Hartigan and Wong, 1979) and the following settings: nstart = 10, iter.max = 100. The optimal number of clusters was determined by Elbow method (Yuan and Yang, 2019). GO analysis of individual clusters was performed using GREAT (4.0.4) (McLean et al., 2010) with single nearest gene setting. Overlap of peaks between clusters was visualized using Intervene(Khan and Mathelier, 2017).

### mRNA-seq data analysis

Raw reads were trimmed by Trim Galore! (0.4.1) with default settings. The trimmed reads were mapped to mm10 reference genome using hisat2 (2.2.1) with default settings. Exon rate was calculated using SeqMonk (1.47.1). Salmon index for decoy-aware mm10 transcriptome was built. After transcript counting, the quant.sf file was processed using Deseq2 (1.36.0)(Love et al., 2014) with custom R script. When correlation between spatial mRNA data and spatial H3K27me3 data was examined, we focused on genes that had substantial level (> 10 counts per gene for RNA) in at least 30% of the slices.

## Acknowledgements

The study was partly supported by United States National Institutes of Health grants R01GM141096 (C.L.) and R01NS123069 (X.J.). The authors also acknowledge support from Institute for Critical Technology and Applied Science at Virginia Tech.

## Author contributions

C.L. conceived and supervised the study. C.L. and D.D.Y. designed the clustering approach. Z.L. conducted ChIP-seq profiling of seizure/control mouse brains and data analysis. C.D. conducted ChIP-seq profiling of normal mouse brains and initial data analysis. Z.Z. conducted RNA-seq profiling of the brain samples. Y.X. contributed to data analysis. S.J. and X.J. produced seizure and control mice. B.Z. helped with the setup of the ChIP-seq experiments. L.B.N. helped with data analysis. Z.L., C.D., and C.L. wrote the manuscript. All authors proofread the manuscript and provided feedback.

## Competing interests

The authors declare no competing interests.

### Data availability

The ChIP-seq and RNA-seq data were deposited into Gene Expression Omnibus under accession number GSE215972.

## REFERENCES

Amini, S., Pushkarev, D., Christiansen, L., Kostem, E., Royce, T., Turk, C., Pignatelli, N., Adey, A., Kitzman, J.O., Vijayan, K., et al. (2014). Haplotype-resolved whole-genome sequencing by contiguity-preserving transposition and combinatorial indexing. Nat Genet 46, 1343–1349. 10.1038/ng.3119.

Bedogni, F., Hodge, R.D., Elsen, G.E., Nelson, B.R., Daza, R.A.M., Beyer, R.P., Bammler, T.K., Rubenstein, J.L.R., and Hevner, R.F. (2010). Tbr1 regulates regional and laminar identity of postmitotic neurons in developing neocortex. Proceedings of the National Academy of Sciences 107, 13129–13134. 10.1073/pnas.1002285107.

Ben-Ari, Y., Tremblay, E., Riche, D., Ghilini, G., and Naquet, R. (1981). Electrographic, clinical and pathological alterations following systemic administration of kainic acid, bicuculline or pentetrazole: metabolic mapping using the deoxyglucose method with special reference to the pathology of epilepsy. Neuroscience 6, 1361–1391. 10.1016/0306-4522(81)90193-7.

Berryer, M.H., Hamdan, F.F., Klitten, L.L., Moller, R.S., Carmant, L., Schwartzentruber, J., Patry, L., Dobrzeniecka, S., Rochefort, D., Neugnot-Cerioli, M., et al. (2013). Mutations in SYNGAP1 cause intellectual disability, autism, and a specific form of epilepsy by inducing haploinsufficiency. Hum Mutat 34, 385–394. 10.1002/humu.22248.

Cao, Z.N., Chen, C.Y., He, B., Tan, K., and Lu, C. (2015). A microfluidic device for epigenomic profiling using 100 cells. Nature Methods 12, 959–962. 10.1038/Nmeth.3488.

Carvill, G.L., Heavin, S.B., Yendle, S.C., McMahon, J.M., O’Roak, B.J., Cook, J., Khan, A., Dorschner, M.O., Weaver, M., Calvert, S., et al. (2013). Targeted resequencing in epileptic encephalopathies identifies de novo mutations in CHD2 and SYNGAP1. Nat Genet 45, 825–830. 10.1038/ng.2646.

Catania, E.H., Pimenta, A., and Levitt, P. (2008). Genetic deletion of Lsamp causes exaggerated behavioral activation in novel environments. Behav Brain Res 188, 380–390. 10.1016/j.bbr.2007.11.022.

Chen, K.H., Boettiger, A.N., Moffitt, J.R., Wang, S., and Zhuang, X. (2015). RNA imaging. Spatially resolved, highly multiplexed RNA profiling in single cells. Science 348, aaa6090. 10.1126/science.aaa6090.

Clement, J.P., Aceti, M., Creson, T.K., Ozkan, E.D., Shi, Y., Reish, N.J., Almonte, A.G., Miller, B. H., Wiltgen, B.J., Miller, C.A., et al. (2012). Pathogenic SYNGAP1 mutations impair cognitive development by disrupting maturation of dendritic spine synapses. Cell 151, 709–723. 10.1016/j.cell.2012.08.045.

de la Fuente Revenga, M., Zhu, B., Guevara, C.A., Naler, L.B., Saunders, J.M., Zhou, Z., Toneatti, R., Sierra, S., Wolstenholme, J.T., Beardsley, P.M., et al. (2021). Prolonged epigenomic and synaptic plasticity alterations following single exposure to a psychedelic in mice. Cell Rep 37, 109836. 10.1016/j.celrep.2021.109836.

Deng, Y., Bartosovic, M., Kukanja, P., Zhang, D., Liu, Y., Su, G., Enninful, A., Bai, Z., Castelo-Branco, G., and Fan, R. (2022a). Spatial-CUT&Tag: Spatially resolved chromatin modification profiling at the cellular level. Science 375, 681–686. 10.1126/science.abg7216.

Deng, Y., Bartosovic, M., Ma, S., Zhang, D., Kukanja, P., Xiao, Y., Su, G., Liu, Y., Qin, X., Rosoklija, G.B., et al. (2022b). Spatial profiling of chromatin accessibility in mouse and human tissues. Nature 609, 375–383. 10.1038/s41586-022-05094-1.

Eng, C.L., Lawson, M., Zhu, Q., Dries, R., Koulena, N., Takei, Y., Yun, J., Cronin, C., Karp, C., Yuan, G.C., and Cai, L. (2019). Transcriptome-scale super-resolved imaging in tissues by RNA seqFISH. Nature 568, 235–239. 10.1038/s41586-019-1049-y.

Feng, H., Khalil, S., Neubig, R.R., and Sidiropoulos, C. (2018). A mechanistic review on GNAO1-associated movement disorder. Neurobiol Dis 116, 131–141. 10.1016/j.nbd.2018.05.005.

Feng, L., and Cooper, J.A. (2009). Dual functions of Dab1 during brain development. Mol Cell Biol 29, 324–332. 10.1128/MCB.00663-08.

Francks, C., Maegawa, S., Lauren, J., Abrahams, B.S., Velayos-Baeza, A., Medland, S.E., Colella, S., Groszer, M., McAuley, E.Z., Caffrey, T.M., et al. (2007). LRRTM1 on chromosome 2p12 is a maternally suppressed gene that is associated paternally with handedness and schizophrenia. Mol Psychiatry 12, 1129–1139, 1057. 10.1038/sj.mp.4002053.

Gonzalez, O.C., Krishnan, G.P., Timofeev, I., and Bazhenov, M. (2019). Ionic and synaptic mechanisms of seizure generation and epileptogenesis. Neurobiol Dis 130, 104485. 10.1016/j.nbd.2019.104485.

Gorkin, D.U., Barozzi, I., Zhao, Y., Zhang, Y., Huang, H., Lee, A.Y., Li, B., Chiou, J., Wildberg, A., Ding, B., et al. (2020). An atlas of dynamic chromatin landscapes in mouse fetal development. Nature 583, 744–751. 10.1038/s41586-020-2093-3.

Guo, H., Bettella, E., Marcogliese, P.C., Zhao, R., Andrews, J.C., Nowakowski, T.J., Gillentine, M.A., Hoekzema, K., Wang, T., Wu, H., et al. (2019). Disruptive mutations in TANC2 define a neurodevelopmental syndrome associated with psychiatric disorders. Nature Communications 10, 4679. 10.1038/s41467-019-12435-8.

Hartigan, J.A., and Wong, M.A. (1979). Algorithm AS 136: A K-Means Clustering Algorithm. Journal of the Royal Statistical Society. Series C (Applied Statistics) 28, 100–108. 10.2307/2346830.

Hauberg, M.E., Creus-Muncunill, J., Bendl, J., Kozlenkov, A., Zeng, B., Corwin, C., Chowdhury, S., Kranz, H., Hurd, Y.L., Wegner, M., et al. (2020). Common schizophrenia risk variants are enriched in open chromatin regions of human glutamatergic neurons. Nat Commun 11, 5581. 10.1038/s41467-020-19319-2.

Heng, J.I.T., Viti, L., Pugh, K., Marshall, O.J., and Agostino, M. (2022). Understanding the impact of ZBTB18 missense variation on transcription factor function in neurodevelopment and disease. Journal of Neurochemistry 161, 219–235. https://doi.org/10.1111/jnc.15572.

Hoxha, E., Marcinno, A., Montarolo, F., Masante, L., Balbo, I., Ravera, F., Laezza, F., and Tempia, F. (2019). Emerging roles of Fgf14 in behavioral control. Behav Brain Res 356, 257–265. 10.1016/j.bbr.2018.08.034.

Inoue, T., Ota, M., Ogawa, M., Mikoshiba, K., and Aruga, J. (2007). Zic1 and Zic3 Regulate Medial Forebrain Development through Expansion of Neuronal Progenitors. The Journal of Neuroscience 27, 5461. 10.1523/JNEUROSCI.4046-06.2007.

Jiang, Y., Matevossian, A., Huang, H.S., Straubhaar, J., and Akbarian, S. (2008). Isolation of neuronal chromatin from brain tissue. BMC Neurosci 9, 42. 10.1186/1471-2202-9-42.

Junker, J.P., Noel, E.S., Guryev, V., Peterson, K.A., Shah, G., Huisken, J., McMahon, A.P., Berezikov, E., Bakkers, J., and van Oudenaarden, A. (2014). Genome-wide RNA Tomography in the zebrafish embryo. Cell 159, 662–675. 10.1016/j.cell.2014.09.038.

Keller, R., Basta, R., Salerno, L., and Elia, M. (2017). Autism, epilepsy, and synaptopathies: a not rare association. Neurol Sci 38, 1353–1361. 10.1007/s10072-017-2974-x.

Khan, A., and Mathelier, A. (2017). Intervene: a tool for intersection and visualization of multiple gene or genomic region sets. BMC Bioinformatics 18, 287. 10.1186/s12859-017-1708-7.

Kim, S.-G., Lee, S., Kim, Y., Park, J., Woo, D., Kim, D., Li, Y., Shin, W., Kang, H., Yook, C., et al. (2021). Tanc2-mediated mTOR inhibition balances mTORC1/2 signaling in the developing mouse brain and human neurons. Nature Communications 12, 2695. 10.1038/s41467-021-22908-4.

Lee, J.H., Daugharthy, E.R., Scheiman, J., Kalhor, R., Yang, J.L., Ferrante, T.C., Terry, R., Jeanty, S.S., Li, C., Amamoto, R., et al. (2014). Highly multiplexed subcellular RNA sequencing in situ. Science 343, 1360–1363. 10.1126/science.1250212.

Lesca, G., Rudolf, G., Bruneau, N., Lozovaya, N., Labalme, A., Boutry-Kryza, N., Salmi, M., Tsintsadze, T., Addis, L., Motte, J., et al. (2013). GRIN2A mutations in acquired epileptic aphasia and related childhood focal epilepsies and encephalopathies with speech and language dysfunction. Nat Genet 45, 1061–1066. 10.1038/ng.2726.

Levesque, M., and Avoli, M. (2013). The kainic acid model of temporal lobe epilepsy. Neurosci Biobehav Rev 37, 2887–2899. 10.1016/j.neubiorev.2013.10.011.

Levesque, M., Langlois, J.M., Lema, P., Courtemanche, R., Bilodeau, G.A., and Carmant, L. (2009). Synchronized gamma oscillations (30-50 Hz) in the amygdalo-hippocampal network in relation with seizure propagation and severity. Neurobiol Dis 35, 209–218. 10.1016/j.nbd.2009.04.011.

Li, H., Handsaker, B., Wysoker, A., Fennell, T., Ruan, J., Homer, N., Marth, G., Abecasis, G., Durbin, R., and Genome Project Data Processing, S. (2009). The Sequence Alignment/Map format and SAMtools. Bioinformatics 25, 2078–2079. 10.1093/bioinformatics/btp352.

Lin, A.Y., Henry, S., Reissner, C., Neupert, C., Kenny, C., Missler, M., Beffert, U., and Ho, A. (2019). A rare autism-associated MINT2/APBA2 mutation disrupts neurexin trafficking and synaptic function. Sci Rep 9, 6024. 10.1038/s41598-019-42635-7.

Lister, R., Mukamel, E.A., Nery, J.R., Urich, M., Puddifoot, C.A., Johnson, N.D., Lucero, J., Huang, Y., Dwork, A.J., Schultz, M.D., et al. (2013). Global epigenomic reconfiguration during mammalian brain development. Science 341, 1237905. 10.1126/science.1237905.

Liu, H., Zhou, J., Tian, W., Luo, C., Bartlett, A., Aldridge, A., Lucero, J., Osteen, J.K., Nery, J.R., Chen, H., et al. (2021). DNA methylation atlas of the mouse brain at single-cell resolution. Nature 598, 120–128. 10.1038/s41586-020-03182-8.

Liu, Y., Yang, M., Deng, Y., Su, G., Enninful, A., Guo, C.C., Tebaldi, T., Zhang, D., Kim, D., Bai, Z., et al. (2020). High-Spatial-Resolution Multi-Omics Sequencing via Deterministic Barcoding in Tissue. Cell 183, 1665–1681 e1618. 10.1016/j.cell.2020.10.026.

Liu, Z., Naler, L.B., Zhu, Y., Deng, C., Zhang, Q., Zhu, B., Zhou, Z., Sarma, M., Murray, A., Xie, H., and Lu, C. (2022). nMOWChIP-seq: low-input genome-wide mapping of non-histone targets. NAR Genom Bioinform 4, lqac030. 10.1093/nargab/lqac030.

Love, M.I., Huber, W., and Anders, S. (2014). Moderated estimation of fold change and dispersion for RNA-seq data with DESeq2. Genome Biol 15, 550. 10.1186/s13059-014-0550-8.

Lu, S., Jang, H., Gu, S., Zhang, J., and Nussinov, R. (2016). Drugging Ras GTPase: a comprehensive mechanistic and signaling structural view. Chemical Society Reviews 45, 4929–4952. 10.1039/C5CS00911A.

Lubeck, E., Coskun, A.F., Zhiyentayev, T., Ahmad, M., and Cai, L. (2014). Single-cell in situ RNA profiling by sequential hybridization. Nat Methods 11, 360–361. 10.1038/nmeth.2892.

Ma, S., Hsieh, Y., Ma, J., and Lu, C. (2018). Low-input and multiplexed microfluidic assay reveals epigenomic variation across cerebellum and prefrontal cortex. Sci Adv 4, eaar8187.

McLean, C.Y., Bristor, D., Hiller, M., Clarke, S.L., Schaar, B.T., Lowe, C.B., Wenger, A.M., and Bejerano, G. (2010). GREAT improves functional interpretation of cis-regulatory regions. Nat Biotechnol 28, 495–501. 10.1038/nbt.1630.

Medvedev, A., Mackenzie, L., Hiscock, J.J., and Willoughby, J.O. (2000). Kainic acid induces distinct types of epileptiform discharge with differential involvement of hippocampus and neocortex. Brain Res Bull 52, 89–98. 10.1016/s0361-9230(00)00239-2.

Möller, C., van Dijk, R.M., Wolf, F., Keck, M., Schönhoff, K., Bierling, V., and Potschka, H. (2019). Impact of repeated kindled seizures on heart rate rhythms, heart rate variability, and locomotor activity in rats. Epilepsy & Behavior 92, 36–44. https://doi.org/10.1016/j.yebeh.2018.11.034.

Murphy, P., and Burnham, W.M. (2003). The effect of kindled seizures on the locomotory behavior of long–evans rats. Experimental Neurology 180, 88–92. https://doi.org/10.1016/S0014-4886(02)00056-0.

Nakamura, K., Kodera, H., Akita, T., Shiina, M., Kato, M., Hoshino, H., Terashima, H., Osaka, H., Nakamura, S., Tohyama, J., et al. (2013). De Novo mutations in GNAO1, encoding a Galphao subunit of heterotrimeric G proteins, cause epileptic encephalopathy. Am J Hum Genet 93, 496–505. 10.1016/j.ajhg.2013.07.014.

Nyegaard, M., Demontis, D., Foldager, L., Hedemand, A., Flint, T.J., Sorensen, K.M., Andersen, P.S., Nordentoft, M., Werge, T., Pedersen, C.B., et al. (2010). CACNA1C (rs1006737) is associated with schizophrenia. Mol Psychiatry 15, 119–121. 10.1038/mp.2009.69.

Picelli, S., Bjorklund, A.K., Reinius, B., Sagasser, S., Winberg, G., and Sandberg, R. (2014a). Tn5 transposase and tagmentation procedures for massively scaled sequencing projects. Genome Res 24, 2033–2040. 10.1101/gr.177881.114.

Picelli, S., Faridani, O.R., Bjorklund, A.K., Winberg, G., Sagasser, S., and Sandberg, R. (2014b). Full-length RNA-seq from single cells using Smart-seq2. Nat Protoc 9, 171–181. 10.1038/nprot.2014.006.

The principles of nerve cell communication. (1997).

Puttachary, S., Sharma, S., Tse, K., Beamer, E., Sexton, A., Crutison, J., and Thippeswamy, T. (2015). Immediate Epileptogenesis after Kainate-Induced Status Epilepticus in C57BL/6J Mice: Evidence from Long Term Continuous Video-EEG Telemetry. PLoS One 10, e0131705. 10.1371/journal.pone.0131705.

Quinlan, A.R., and Hall, I.M. (2010). BEDTools: a flexible suite of utilities for comparing genomic features. Bioinformatics 26, 841–842. 10.1093/bioinformatics/btq033.

Raimondo, J.V., Burman, R.J., Katz, A.A., and Akerman, C.J. (2015). Ion dynamics during seizures. Front Cell Neurosci 9, 419. 10.3389/fncel.2015.00419.

Rakic, P., Ayoub, A.E., Breunig, J.J., and Dominguez, M.H. (2009). Decision by division: making cortical maps. Trends Neurosci 32, 291–301. 10.1016/j.tins.2009.01.007.

Rodriques, S.G., Stickels, R.R., Goeva, A., Martin, C.A., Murray, E., Vanderburg, C.R., Welch, J., Chen, L.M., Chen, F., and Macosko, E.Z. (2019). Slide-seq: A scalable technology for measuring genome-wide expression at high spatial resolution. Science 363, 1463–1467. 10.1126/science.aaw1219.

Sharma, S., Puttachary, S., Thippeswamy, A., Kanthasamy, A.G., and Thippeswamy, T. (2018). Status Epilepticus: Behavioral and Electroencephalography Seizure Correlates in Kainate Experimental Models. Front Neurol 9, 7. 10.3389/fneur.2018.00007.

Shaw-Smith, C., Pittman, A.M., Willatt, L., Martin, H., Rickman, L., Gribble, S., Curley, R., Cumming, S., Dunn, C., Kalaitzopoulos, D., et al. (2006). Microdeletion encompassing MAPT at chromosome 17q21.3 is associated with developmental delay and learning disability. Nat Genet 38, 1032–1037. 10.1038/ng1858.

Singh, K., Loreth, D., Pottker, B., Hefti, K., Innos, J., Schwald, K., Hengstler, H., Menzel, L., Sommer, C.J., Radyushkin, K., et al. (2018). Neuronal Growth and Behavioral Alterations in Mice Deficient for the Psychiatric Disease-Associated Negr1 Gene. Front Mol Neurosci 11, 30. 10.3389/fnmol.2018.00030.

Smits, J.J., Oostrik, J., Beynon, A.J., Kant, S.G., de Koning Gans, P.A.M., Rotteveel, L.J.C., Klein Wassink-Ruiter, J.S., Free, R.H., Maas, S.M., van de Kamp, J., et al. (2019). De novo and inherited loss-of-function variants of ATP2B2 are associated with rapidly progressive hearing impairment. Hum Genet 138, 61–72. 10.1007/s00439-018-1965-1.

Stewart, L.S., and Leung, L.S. (2003). Temporal lobe seizures alter the amplitude and timing of rat behavioral rhythms. Epilepsy & Behavior 4, 153–160. https://doi.org/10.1016/S1525-5050(03)00006-4.

Stogmann, E., Reinthaler, E., Eltawil, S., El Etribi, M.A., Hemeda, M., El Nahhas, N., Gaber, A.M., Fouad, A., Edris, S., Benet-Pages, A., et al. (2013). Autosomal recessive cortical myoclonic tremor and epilepsy: association with a mutation in the potassium channel associated gene CNTN2. Brain 136, 1155–1160. 10.1093/brain/awt068.

Strang, K.H., Golde, T.E., and Giasson, B.I. (2019). MAPT mutations, tauopathy, and mechanisms of neurodegeneration. Lab Invest 99, 912–928. 10.1038/s41374-019-0197-x.

Strehlow, V., Heyne, H.O., Vlaskamp, D.R.M., Marwick, K.F.M., Rudolf, G., de Bellescize, J., Biskup, S., Brilstra, E.H., Brouwer, O.F., Callenbach, P.M.C., et al. (2019). GRIN2A-related disorders: genotype and functional consequence predict phenotype. Brain 142, 80–92. 10.1093/brain/awy304.

Tadmouri, A., Kiyonaka, S., Barbado, M., Rousset, M., Fablet, K., Sawamura, S., Bahembera, E., Pernet-Gallay, K., Arnoult, C., Miki, T., et al. (2012). Cacnb4 directly couples electrical activity to gene expression, a process defective in juvenile epilepsy. EMBO J 31, 3730–3744. 10.1038/emboj.2012.226.

Venkateswaran, S., Myers, K.A., Smith, A.C., Beaulieu, C.L., Schwartzentruber, J.A., Consortium, F.C., Majewski, J., Bulman, D., Boycott, K.M., and Dyment, D.A. (2014). Whole-exome sequencing in an individual with severe global developmental delay and intractable epilepsy identifies a novel, de novo GRIN2A mutation. Epilepsia 55, e75–79. 10.1111/epi.12663.

Wang, X., Allen, W.E., Wright, M.A., Sylwestrak, E.L., Samusik, N., Vesuna, S., Evans, K., Liu, C., Ramakrishnan, C., Liu, J., et al. (2018). Three-dimensional intact-tissue sequencing of single-cell transcriptional states. Science 361. 10.1126/science.aat5691.

Yon Rhee, S., Wood, V., Dolinski, K., and Draghici, S. (2008). Use and misuse of the gene ontology annotations. Nature Reviews Genetics 9, 509–515. 10.1038/nrg2363.

Yuan, C., and Yang, H. (2019). Research on K-Value Selection Method of K-Means Clustering Algorithm. J 2, 226–235. 10.3390/j2020016.

Zeng, L.H., Xu, L., Rensing, N.R., Sinatra, P.M., Rothman, S.M., and Wong, M. (2007). Kainate seizures cause acute dendritic injury and actin depolymerization in vivo. J Neurosci 27, 11604–11613. 10.1523/JNEUROSCI.0983-07.2007.

Zhao, T., Chiang, Z.D., Morriss, J.W., LaFave, L.M., Murray, E.M., Del Priore, I., Meli, K., Lareau, C.A., Nadaf, N.M., Li, J., et al. (2022). Spatial genomics enables multi-modal study of clonal heterogeneity in tissues. Nature 601, 85–91. 10.1038/s41586-021-04217-4.

Zhao, Y., and Garcia, B.A. (2015). Comprehensive Catalog of Currently Documented Histone Modifications. Cold Spring Harbor Perspectives in Biology 7. 10.1101/cshperspect.a025064.

Zhu, B., Hsieh, Y.P., Murphy, T.W., Zhang, Q., Naler, L.B., and Lu, C. (2019). MOWChIP-seq for low-input and multiplexed profiling of genome-wide histone modifications. Nat Protoc 14, 3366–3394. 10.1038/s41596-019-0223-x.

