## Supplementary Information for "Epigenomic tomography for probing spatially-defined molecular state in the brain"

### Supplementary Figures

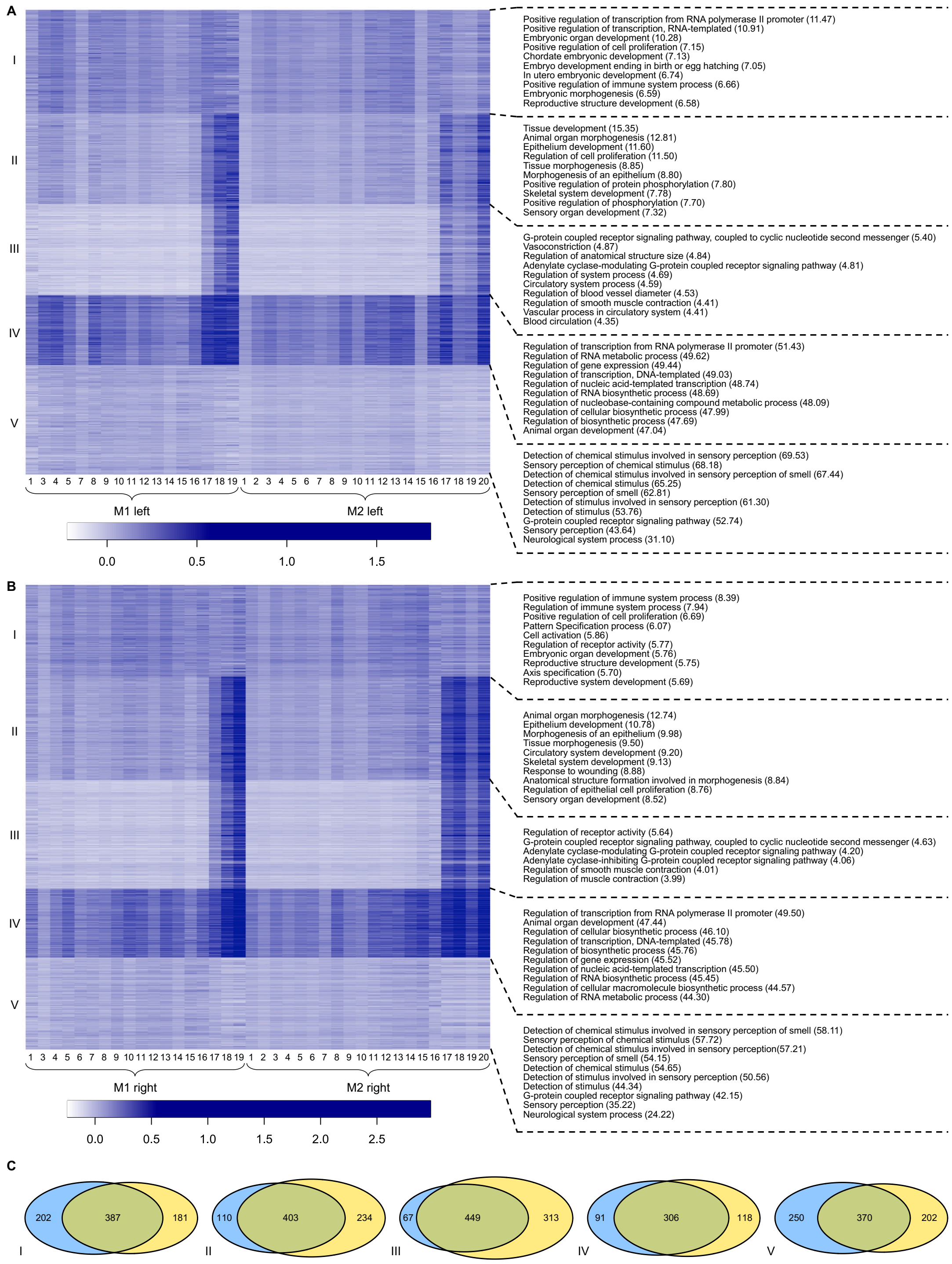


**Figure S1. Double epigenomic tomography generated by k-means clustering of promoter H3K27me3 peaks using neuronal data on neocortex of the same side of different mice (M1 and M2) and top GO biological process terms associated with each cluster.** (A) left neocortex. (B) right neocortex. (C) Overlap of peaks between clusters of the same spatial pattern from the two epigenomic tomographies. Top 10 GO biological process terms were shown if the total number of GO terms is greater than 10. The full list of GO terms is presented in Supplementary Dataset S6.

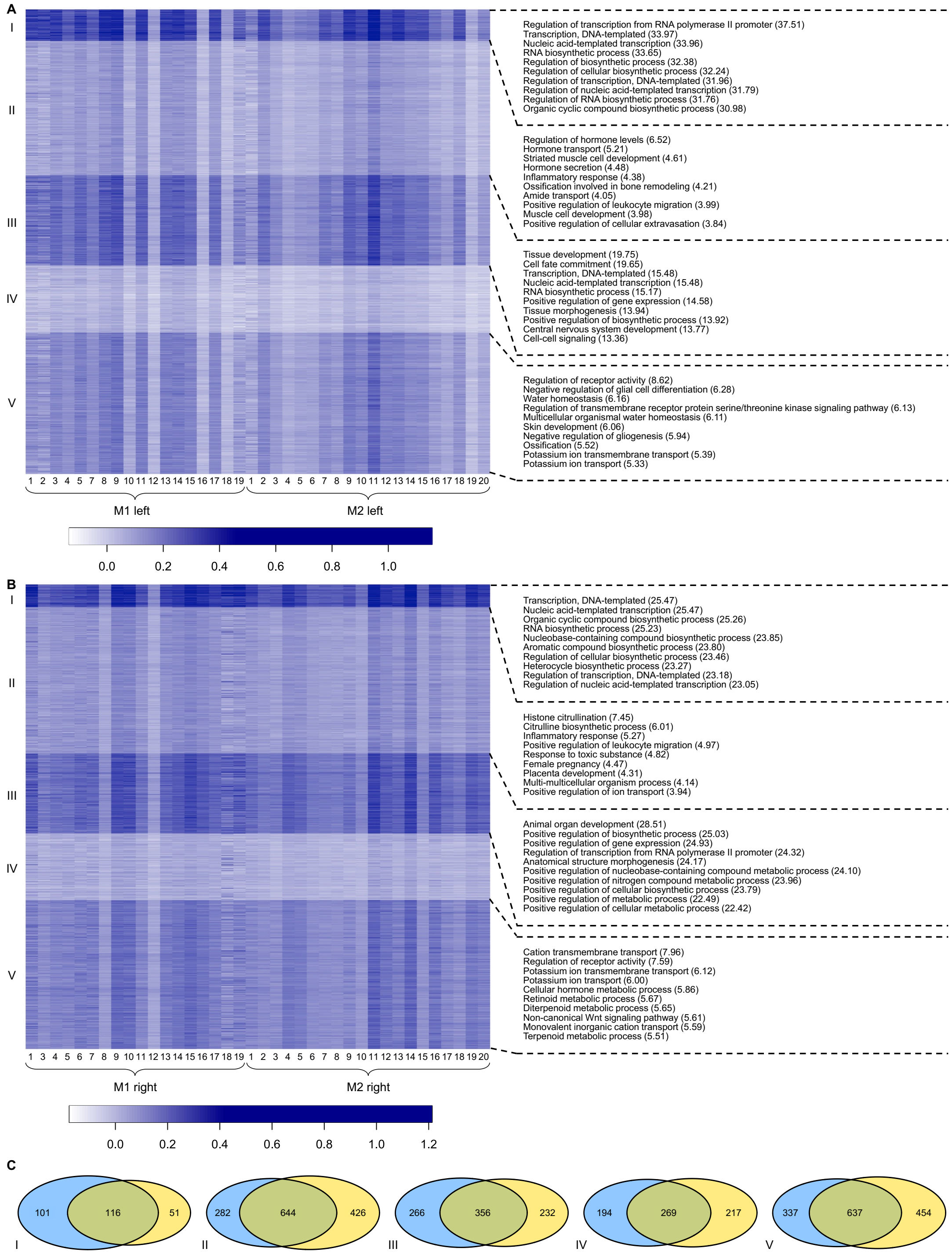


**Figure S2. Double epigenomic tomography generated by k-means clustering of promoter H3K27me3 peaks using glial data on the neocortex of the same side of different mice (M1 and M2) and top GO biological process terms associated with each cluster.** (A) left neocortex. (B) right neocortex. (C) Overlap of peaks between clusters of the same spatial pattern from the two epigenomic tomographies. Top 10 GO biological process terms were shown if the total number of GO terms is greater than 10. The full list of GO terms is presented in Supplementary Dataset S7.

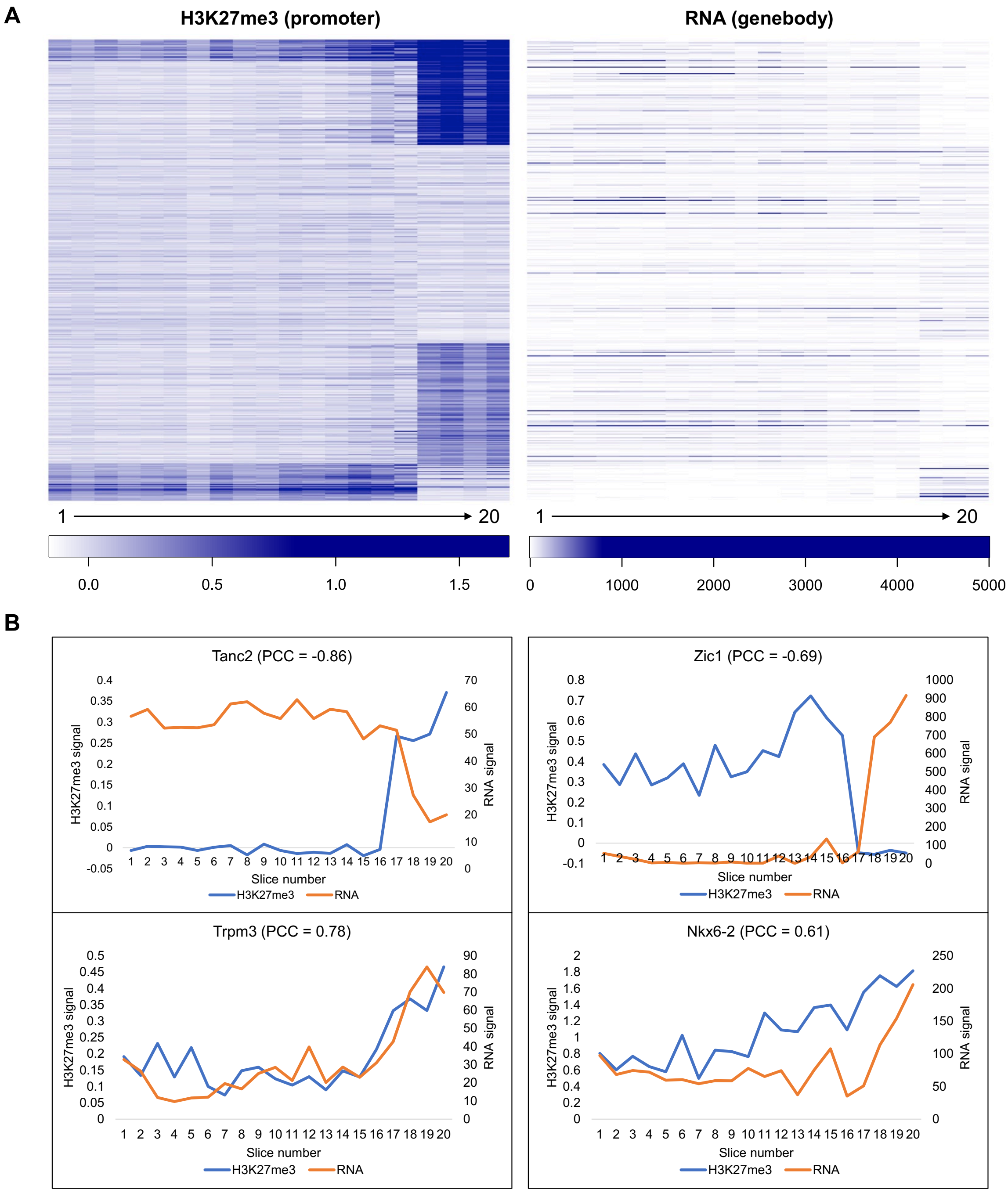


**Figure S3. Correlation between promoter H3K27me3 and genebody RNA signal.** H3K27me3 data on neurons of M2 and RNA-seq data on neurons of M3 are used. (A) k-means clustering of promoter H3K27me3 signal for 495 genes with an inverse trend between H3K27me3 and RNA (Pearson's correlation coefficient or PCC < -0.5). Genebody RNA signal for the genes ranked in the same order are shown on the right. (B) Representative genes with either strong inverse or positive correlation between their promoter H3K27me3 and RNA signal.

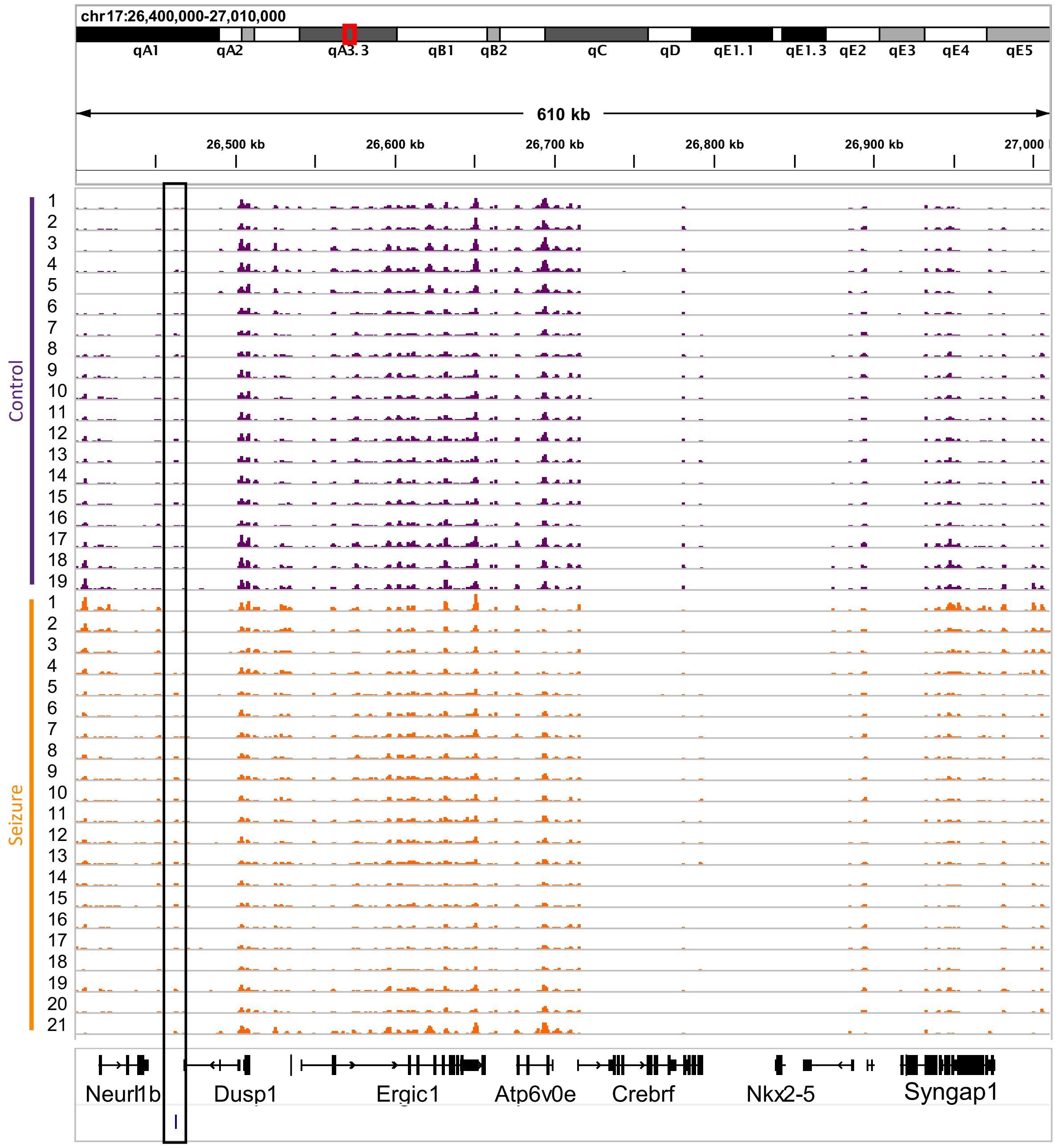


**Figure S4. Normalized H3K27ac signal of neurons from control and seizure mice at *Syngap1*.** The predicted enhancer of *Syngap1* is marked. The enhancer peak was identified by selecting H3K27ac peak within ± 1M bp from TSS of a gene with the highest correlation with the gene expression (mRNA-seq) data.

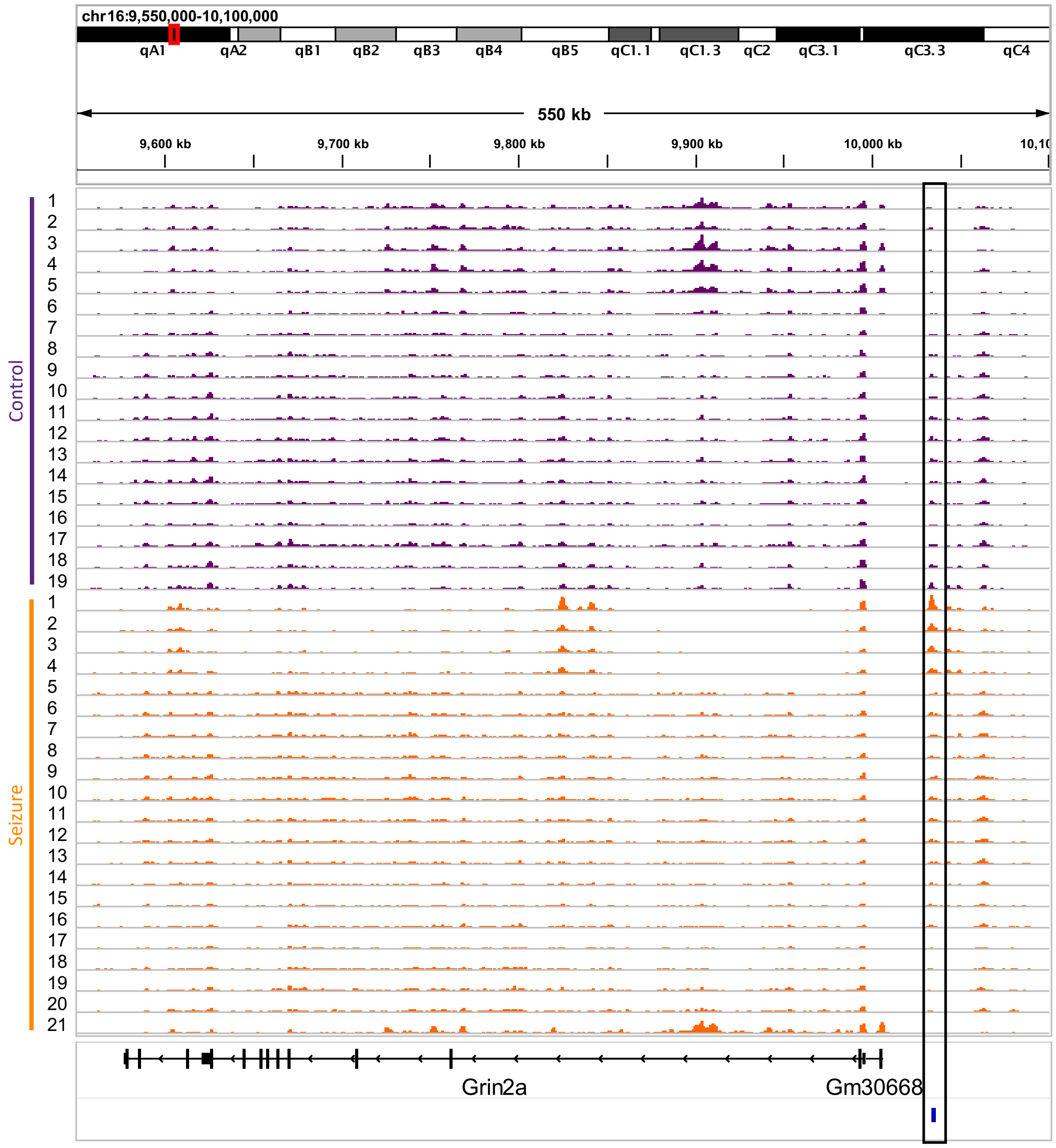


**Figure S5. Normalized H3K27ac signal of neurons from control and seizure mice at *Grin2a*.** The predicted enhancer of *Grin2a* is marked.

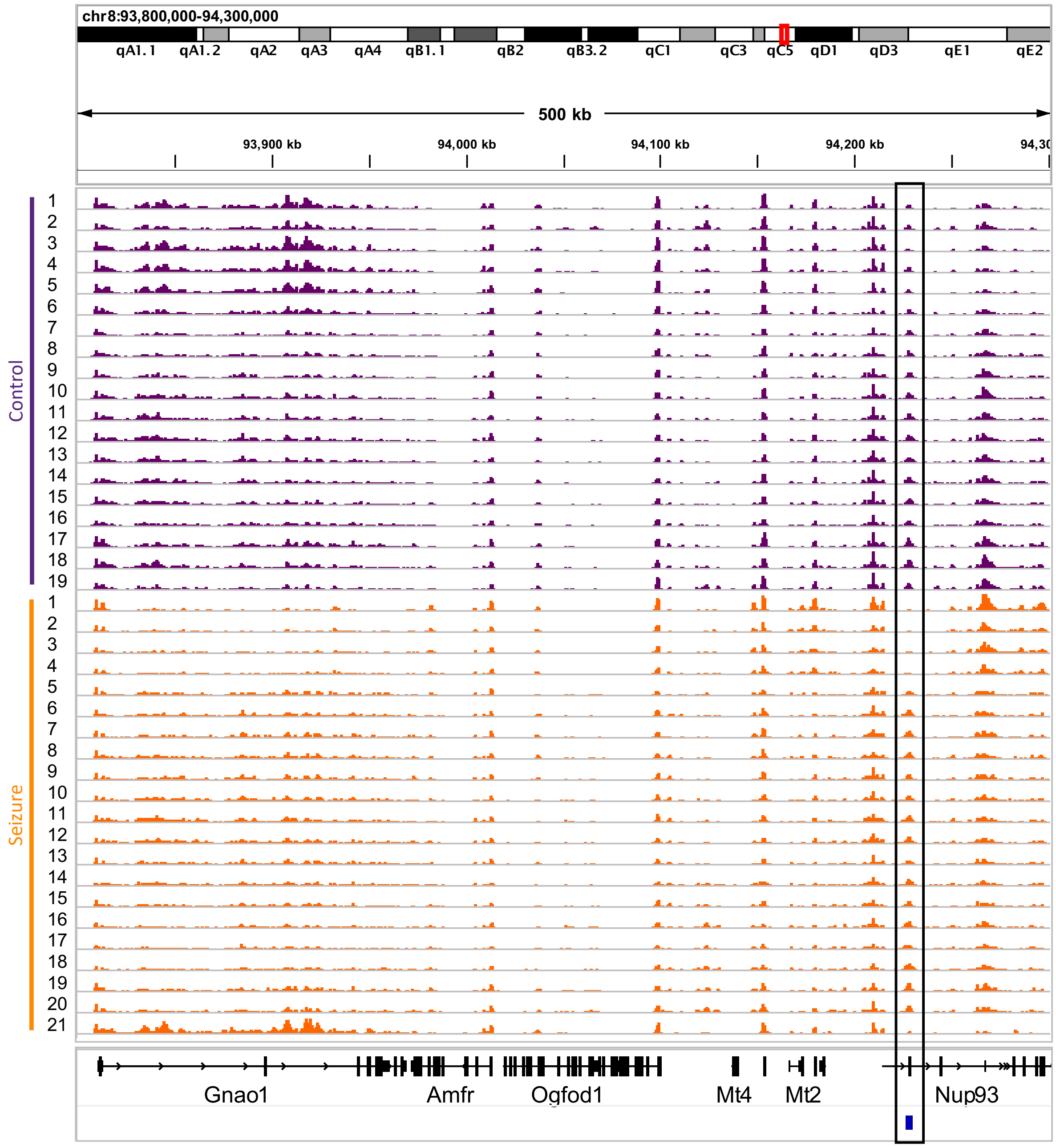


**Figure S6. Normalized H3K27ac signal of neurons from control and seizure mice at *Gnao1*.** The predicted enhancer of *Gnao1* is marked.

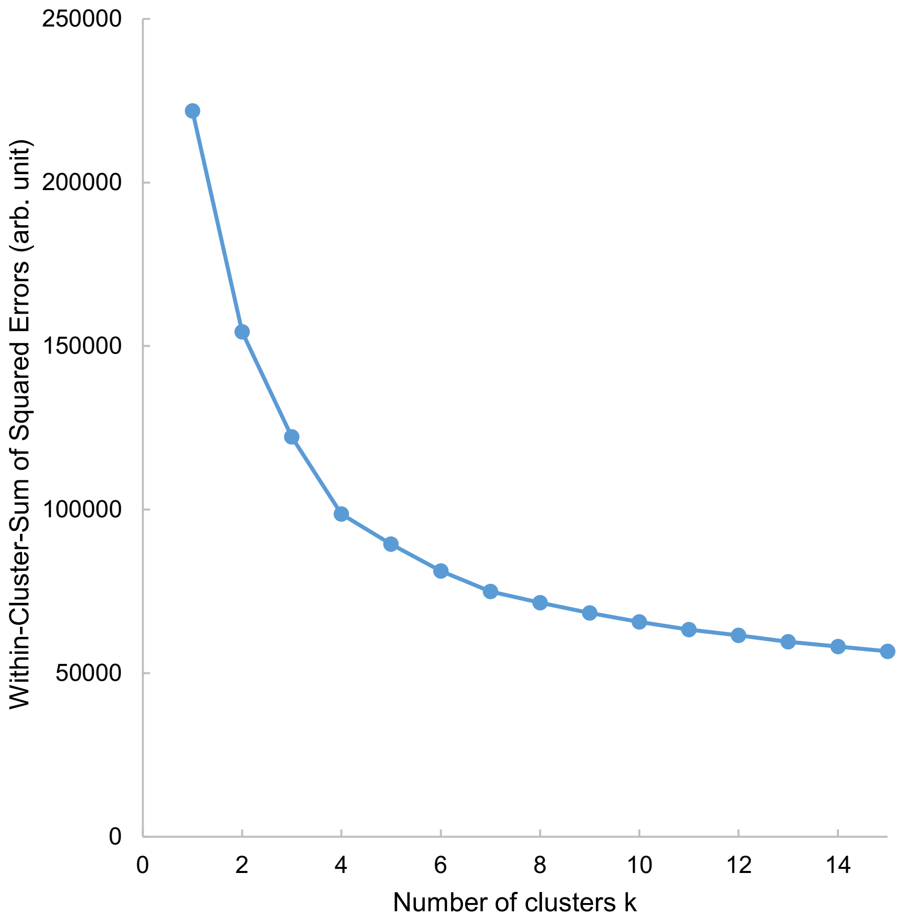


**Figure S7. Elbow method to determine the optimal number of clusters k for k-means clustering of enhancer peaks.** M4-5 right H3K27ac data set was used in the analysis.

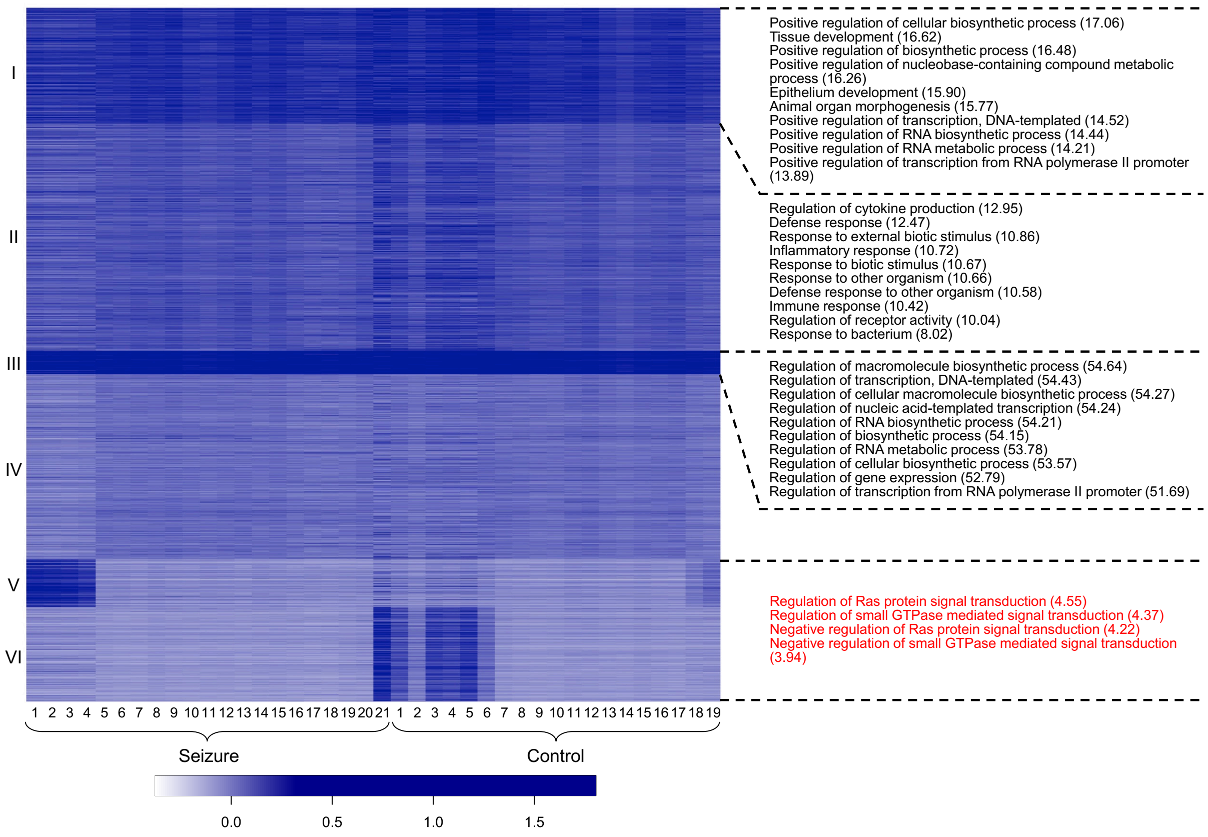


**Figure S8. Double epigenomic tomography generated by k-means clustering of promoter H3K27me3 peaks using neuronal data on the right neocortex of both seizure and control mice and top GO biological process terms associated with each cluster.** k=6. Top 10 GO biological process terms were shown if the total number of GO terms is greater than 10. GO terms associated with differential clusters are in red. The full list of GO terms is presented in Supplementary Dataset S8.

**Supplementary Datasets**
**Supplementary Dataset S1 Complete lists of GO terms associated with the clusters in Figure 2.**
**Supplementary Dataset S2 Complete lists of GO terms associated with the clusters in Figure 3.**
**Supplementary Dataset S3 Complete lists of GO terms associated with the clusters in Figure 4.**
**Supplementary Dataset S4 Complete lists of GO terms associated with the clusters in Figure 5.**
**Supplementary Dataset S5 Complete lists of GO terms associated with the clusters in Figure 6.**
**Supplementary Dataset S6 Complete lists of GO terms associated with the clusters in Supplementary Figure S1.**
**Supplementary Dataset S7 Complete lists of GO terms associated with the clusters in Supplementary Figure S2.**
**Supplementary Dataset S8 Complete lists of GO terms associated with the clusters in Supplementary Figure S8.**

**Supplementary Table S1.** Quality control metrics of MOWChIP-seq datasets on mouse neocortex slices.

**Supplementary Video S1.** Kainic acid-induced seizure. The video shows a mouse with its tail marked with 2 lines (denoting 2 injections of kainic acid) is experiencing seizure. The mouse can be seen jumping and wobbling, then standing up with its front legs twitching.
